# Novel Protoporphyrinogen oxidase 1 mutations endow resistance to PPO-inhibiting herbicides in *Bassia scoparia*

**DOI:** 10.64898/2026.01.15.699694

**Authors:** Aimone Porri, Quincy D. Law, Charles M. Geddes, Joseph Ikley, Samuel Willingham, Philipp Johnen, Ingo Meiners, Sama Al-Sammarraie, Kim Crommar, Pieter B.F. Ouwerkerk, Michael Betz, Brian M. Jenks, Heike Heiser, Fabienne Baumann, Frank Braendle, Jens Lerchl

## Abstract

PPO-inhibiting herbicides are widely used to manage weeds in different cropping systems, yet resistance evolution threatens their long-term efficacy. Here, we investigated the molecular basis of resistance to PPO-inhibiting herbicides in *Bassia scoparia* biotypes collected from four locations in North Dakota, USA. Greenhouse dose–response assays revealed high levels of resistance to saflufenacil and carfentrazone-ethyl, while fomesafen retained full efficacy across all biotypes. Resistant plants did not show increased copy number or elevated expression of *PPO1* or *PPO2*. Sequencing of survivor plants revealed conserved PPO2 sequences, but consistent target-site substitutions at position F454 in PPO1, including F454I, F454L, and F454V. *In vitro* enzyme assays demonstrated that these substitutions impair PPO1 sensitivity to saflufenacil and carfentrazone-ethyl, but not to fomesafen. Ectopic expression of *B. scoparia* PPO1 F454 mutant variants in *Arabidopsis thaliana* conferred tolerance to saflufenacil and carfentrazone-ethyl, but not to fomesafen, supporting greenhouse and *in vitro* results. Molecular modeling indicated that the conformational flexibility and interaction profile of fomesafen enables it to maintain binding to mutated PPO1 variants, in contrast to the more rigid structures of saflufenacil and carfentrazone-ethyl. A yeast-based complementation system further confirmed that F454 substitutions decrease herbicide sensitivity. In addition, developmental profiling showed distinct expression patterns of *PPO1* and *PPO2* during early growth stages in *B. scoparia* and *Amaranthus* spp., highlighting isoform-specific roles. Together, these findings represent the first reported PPO1 target-site mutations in a broadleaf weed species as a key mechanism of resistance and highlight that fomesafen is effective to control resistant *B. scoparia* populations.

## 1. Introduction

Herbicide resistance is a major challenge in chemical weed control, as resistant populations can develop faster than new active ingredients are discovered and commercialized (Gaines et al., 2021). Despite this challenge, reliance on herbicides for weed management continues to increase as urbanization results in dwindling rural populations relocating to urban centers and concomitant decreasing on-farm labor (Peterson et al., 2018). The resultant selection pressure for resistance caused by greater reliance on herbicides and fewer effective herbicide options has led to a steady increase in herbicide-resistant weeds worldwide (Heap, 2025). Stacking of resistance traits in multiple herbicide-resistant weed populations further complicates weed control (Torra et al., 2021), leading to a simultaneous need for new active ingredients to manage them combined with integrated weed management strategies to curb further selection and spread.

*Bassia scoparia* (L.) A.J. Scott (kochia) is a competitive, warm-season (C4) summer-annual tumbleweed that is challenging to manage in croplands and ruderal areas due to its unique biology. Early emergence and prolonged emergence periodicity (Schwinghamer and Van Acker, 2008; Dille et al., 2017; Kumar et al., 2018), tolerance to abiotic stress, high fecundity (Friesen et al., 2009; Geddes and Sharpe, 2022), high genetic diversity (Martin et al., 2020), and efficient pollen- and seed-mediated gene flow (Beckie et al., 2016) contribute to the persistence and spread of this broadleaf weed in arid and semi-arid environments. Recent surveys of the United States and Canada identified *B. scoparia* as the second- and third-most troublesome weed in broadleaf and grass crops, respectively (Van Wychen, 2023, 2025). *Bassia scoparia* was the sixth most-abundant weed present after postemergence weed control among 4077 annual-cropped fields surveyed in the Canadian Prairies, where it has increased rapidly in the past decade (Leeson et al., 2025). Uncontrolled *B. scoparia* plants can cause severe (>90%) yield losses in crops such as corn (*Zea mays* L.), sorghum [*Sorghum bicolor* (L.) Moench ssp. bicolor], sugar beet (*Beta vulgaris* L.), and sunflower (*Helianthus annuus* L.) due to their competitiveness and resource-independent interference (Geddes and Sharpe, 2022). Indeterminate growth often results in lush green *B. scoparia* plants in senesced crops during harvest, and concomitant harvest inefficiencies.

*Bassia scoparia* populations in North America have been reported with resistance to five herbicide sites of action, including auxin mimics, and inhibitors of acetolactate synthase (ALS), photosystem II (PSII), 5-enolpyruvylshikimaate-3-phosphate synthase (EPSPS), and protoporphyrinogen oxidase (PPO) [Herbicide Resistance Action Committee (HRAC) Groups 4, 2, 5, 9, and 14, respectively] (Heap, 2025). Resistance to ALS- and EPSPS-inhibiting herbicides is widespread among *B. scoparia* populations in the North American Great Plains, while auxin mimic and PSII inhibitor resistance occur less frequently and vary geographically within this region (Geddes et al., 2022; Geddes et al., 2023; Sharpe et al., 2023; Araujo et al., 2024; Dhanda et al., 2025). *Bassia scoparia* populations with resistance to these four sites of action have been known to occur for over a decade (Kumar et al., 2019). In contrast, PPO inhibitor resistance was confirmed only recently in *B. scoparia* populations collected after control failures in Saskatchewan, Canada in 2021, and two locations in North Dakota, USA in 2022 (Geddes et al., 2025a). All three populations were highly resistant to foliar-applied saflufenacil and carfentrazone-ethyl, while other PPO-inhibiting herbicides were not tested. Screening of *B. scoparia* survey (n = 882) and grower-submitted (n = 14) samples in Western Canada identified an additional PPO inhibitor-resistant population in Alberta, Canada in 2023 (Geddes et al., 2025b).

PPO–inhibiting herbicides are used across agronomic and specialty crops to control broadleaf and some grass weeds through pre-plant burndown, preemergence residual, and postemergence applications (Barker et al., 2023). These herbicides function by inhibiting the PPO enzyme, a key component in the chlorophyll biosynthesis pathway, leading to a rapid accumulation of phytotoxic intermediates that cause cell membrane disruption and plant death. Adoption of PPO-inhibiting herbicides has increased in recent years, largely driven by the ongoing evolution and spread of ALS and EPSPS inhibitor-resistant weeds (Dayan et al., 2018; Barker et al., 2023). Use may continue to expand, as PPO represents a versatile enzymatic target, with more than 100,000 experimental molecules identified as potential inhibitors. Previous research explored combining structural features from commercial PPO inhibitors as a strategy to overcome PPO inhibitor resistance (Mattison et al., 2023).

Protoporphyrinogen oxidase catalyzes chlorophyll and heme production within chloroplasts (Barker et al., 2023). The two organelle-specific isoforms of PPO (PPO1 and PPO2) are encoded by two nuclear genes (PPX1 and PPX2, respectively). To-date, most cases of target-site PPO inhibitor resistance have been linked to mutations in PPX2 (Dayan et al., 2018; Barker et al., 2023; Riechers et al., 2024). The PPX2 gene appears to provide a more evolutionarily favorable pathway for the development of resistance because PPO2 was initially reported to be targeted to both chloroplasts and mitochondria in species such as *Spinacia oleracea* (Watanabe et al., 2001), *Amaranthus tuberculatus*, and *A. palmeri* (Dayan et al., 2018). However, more recent studies indicate that in *Arabidopsis, Spinacia and Amaranthus* PPO1 and PPO2 are both localized in the chloroplast (Hedtke et al., 2023; Wittmann et al., 2024). Further, Dayan et al. (2018) identified *B. scoparia* as a species with the same PPO2 tandem repeat facilitating the Gly210 deletion (ΔG210) that confers resistance to PPO inhibitors in other members of the Amaranthaceae family, *A. tuberculatus* and *A. palmeri*.

### Objectives

While the presence of PPO inhibitor resistance in *B. scoparia* has been confirmed (Geddes et al., 2025a), the specific molecular and biochemical mechanisms driving it have not been elucidated. Thus, the objectives of this research were to: 1) confirm resistance to saflufenacil, carfentrazone-ethyl, and fomesafen in four North Dakota biotypes; 2) investigate the roles of target-site mutations, gene amplification, and gene overexpression as potential resistance mechanisms; and 3) functionally validate the impact of identified mutations on herbicide sensitivity.

## 2. Materials and methods

### Plant material and greenhouse herbicidal treatments

A greenhouse experiment was conducted to evaluate whole-plant response to foliar applications of saflufenacil, carfentrazone-ethyl, and fomesafen, on four *B. scoparia* biotypes collected from field populations near Minot, Mandan, Mott and Berthold, North Dakota in 2023. Additionally, a known sensitive biotype was included for comparison.

For each biotype, the experiment was designed as a three-factor (biotype x herbicide x herbicide rate) factorial on a randomized complete block design (RCBD), with 12 replications, and repeated once. Seeds from each biotype were sown in greenhouse flats measuring 25 by 50 cm, containing commercial potting mix (Fafard Germinating Mix; Sun Gro Horticulture, Agawa, MA), and transplanted into 760-cm^3^ pots (Ray Leach SC-10 Super Cell Cone-tainers; Stuewe & Sons, Tangent, OR) filled with the same potting mix as sown once seedlings reached 3 cm in height. Plants were watered daily and fertilized weekly with a micro- and macronutrient fertilizer (Jack’s Classic Professional 20-20-20, JR Peters Inc., Allentown PA) until they reached the 7 to 10 cm in height, at which time herbicide applications were made using a track-mounted research sprayer (Generation III Research Sprayer, DeVries Manufacturing, Hollandale, MN) calibrated to deliver 140 L ha^-1^ at 207 kPa via an Turbo TeeJet^®^ TT110015 nozzle (TeeJet Technologies, Glendale Heights, IL). Herbicide treatments included seven rates of carfentrazone-ethyl (0 to 210 g ai ha^-1^, projected labeled 1X use rate = 17.5 g ai ha^-1^), saflufenacil (0 to 300 g ai ha^-1^, labeled 1X use rate = 25 g ai ha^-1^), or fomesafen (0 to 2530 g ai ha^-1^, labeled 1X use rate = 210 g ai ha^-1^) with methylated seed oil (MSO Ultra, Precision Laboratories, Waukegan, IL) added to each herbicide treatment at 1% v v^-1^. Treatments were applied at 0.5X, 1X, 2X, 4X, 6X, 8X and 12X of the recommended field use rate. Greenhouse environmental conditions included a 14:10 h light:dark photoperiod, where natural light was supplemented with high-pressure sodium bulbs delivering 1100 μmol m^-2^ s^-1^ photon flux during daylight hours, and day/night temperatures of 30 and 25°C, respectively. Visual estimates of *B. scoparia* control were recorded at 7, 14 and 21 days after application (DAA) utilizing a 0 to 100 scale, where 0 = no injury and 100 = complete plant death. Data were subjected to ANOVA statistical analysis.

### Yeast assay

A haploid strain YKC027 (MATα his3Δ1 leu2Δ0 ura3Δ0 lys2Δ0 hem14::kanMX4) knock-out for the single PPO gene HEM14 (Glerum et al., 1996) was isolated by tetrad dissection and mass-sporulation of strain YER014 BY4743 which is heterozygous for HEM14/hem14::kanMX4 and which originates from the yeast genome deletion collection (Transomic, AL, USA). Haploids were selected on G418 containing medium thereby selecting for the hem14::kanMX knock-out locus which were genotyped with the ABCD primer system (Cherry et al., 2011; Giaever and Nislow, 2014). The hem14::kanMX haploid strain YKC027 was further genotyped and phenotyped by selection on minimal SC plates to establish phenotypes for lysine and methionine auxotrophy versus prototrophy. Constructs p416ADH-AmPPO1 and p416ADH-AmPPO2 were based on *A. palmeri* PPO1 and PPO2 sequences, respectively. The mating type of YKC027 was confirmed with colony PCR using the primers described by Huxley et al. (1990). Although YKC027 lacks mitochondrial respiration because no active PPO gene is present, it will still grow on standard YPD medium containing glucose as fermentable carbon source. If glucose is replaced by 2% ethanol and 3% glycerol as non-fermentable carbon sources in YPGE medium, YKC027 cannot grow anymore since yeast needs mitochondria with a fully functional and heme-dependent respiration system to be able to grow on glycerol and ethanol. Growth on YPGE can be restored again by transforming YKC027 with a plant PPO gene construct. With the conditional lethality screening system that we created in this way, we were able to use a simple plate-based screening system to check for herbicide sensitivity versus tolerance conferred by plant PPO genes or PPO variants since inhibition of PPO activity by an herbicide will inhibit growth when cultured on YPGE. Plant PPO constructs for complementation of the hem14::kanMX4 locus in YKC027 were synthetically made (Azenta-Genewiz, Germany) in the URA3 ARS-CEN vector p416ADH in which the gene of interest is driven by the ADH1 promoter (Mumberg et al., 1996), transformed into yeast by a LiAc transformation method (Gietz and Schiestl, 2008) and selected for uracil prototrophy on SC (Synthetic Complete) 2% glucose plates respecting the genotype from Table 1. Apart from the wild-type BsPPO1 sequence from *B. scoparia* (represented by construct p416ADH-BsPP01 in yeast strain YSA024), three different point mutations were analyzed at position F454 (Table 1, Yeast strains YSA025 to YSA027 harboring constructs p416ADH-BsPP01-F454I, p416ADH-BsPP01-F454V, p416ADH-BsPP01-F454L).. For herbicide screens, colonies were grown on YPGE plates with or without PPO-inhibiting herbicides to check for tolerance levels. The PPO inhibiting herbicides saflufenacil, carfentrazone and fomesafen were prepared as 0.1 M DMSO stocks and from there, diluted to 1 mM and 10 mM stocks, also in DMSO, to serve as 1000-fold stocks for the herbicide growth assays. The integrity of the PPO complementation constructs in yeast was checked by colony PCR and sequencing (LGC Genomics or Plasmidsaurus) using primers ProADH Fw1 GCTATCAAGTATAAATAGACCTG and Tcyc1 Rev1 GTATAATGTTACATGCGTACAC.

**Table 1.**
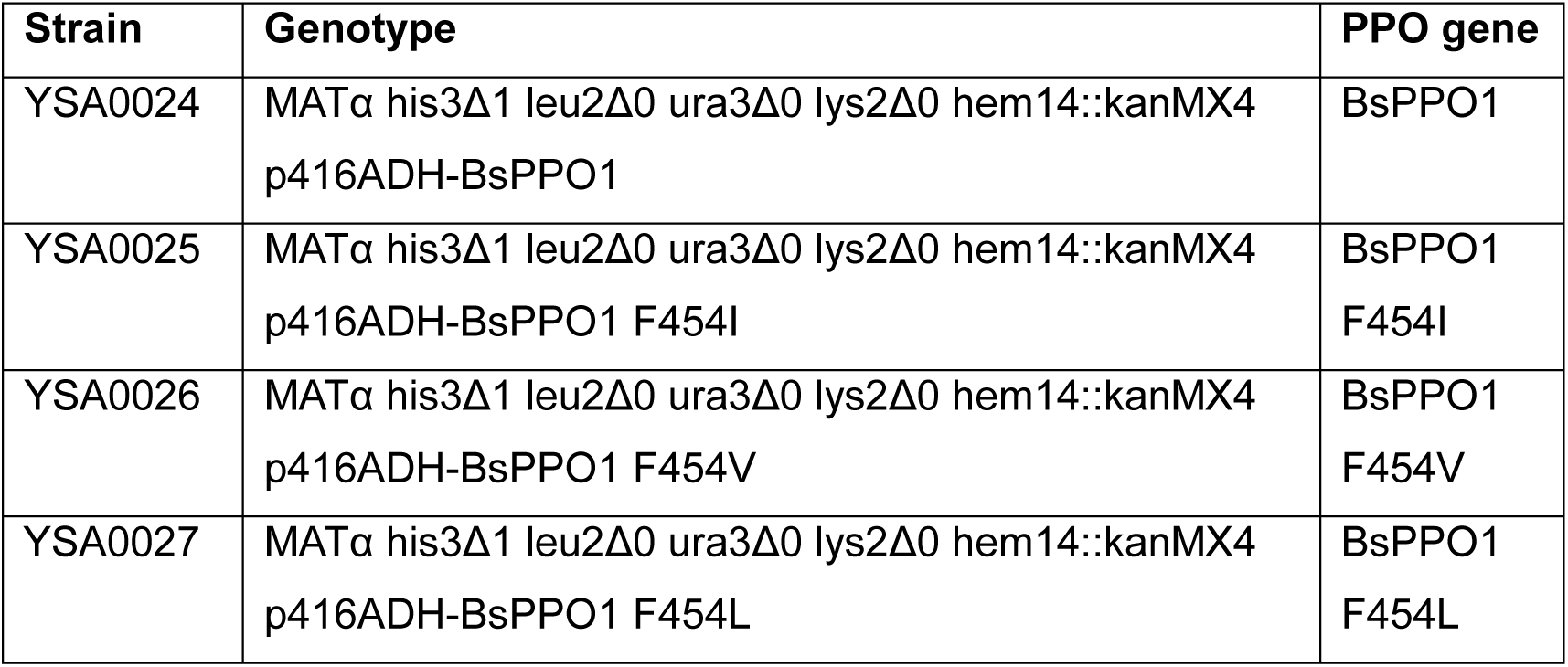
Overview of yeast strains with or without PPO complementation constructs used in this study.

| Strain | Genotype | PPO gene |
| --- | --- | --- |
| YSA0024 | MATα his3Δ1 leu2Δ0 ura3Δ0 lys2Δ0 hem14::kanMX4<br>p416ADH-BsPPO1 | BsPPO1 |
| YSA0025 | MATα his3Δ1 leu2Δ0 ura3Δ0 lys2Δ0 hem14::kanMX4<br>p416ADH-BsPPO1 F454I | BsPPO1<br>F454I |
| YSA0026 | MATα his3Δ1 leu2Δ0 ura3Δ0 lys2Δ0 hem14::kanMX4<br>p416ADH-BsPPO1 F454V | BsPPO1<br>F454V |
| YSA0027 | MATα his3Δ1 leu2Δ0 ura3Δ0 lys2Δ0 hem14::kanMX4<br>p416ADH-BsPPO1 F454L | BsPPO1<br>F454L |

**Table 1.**
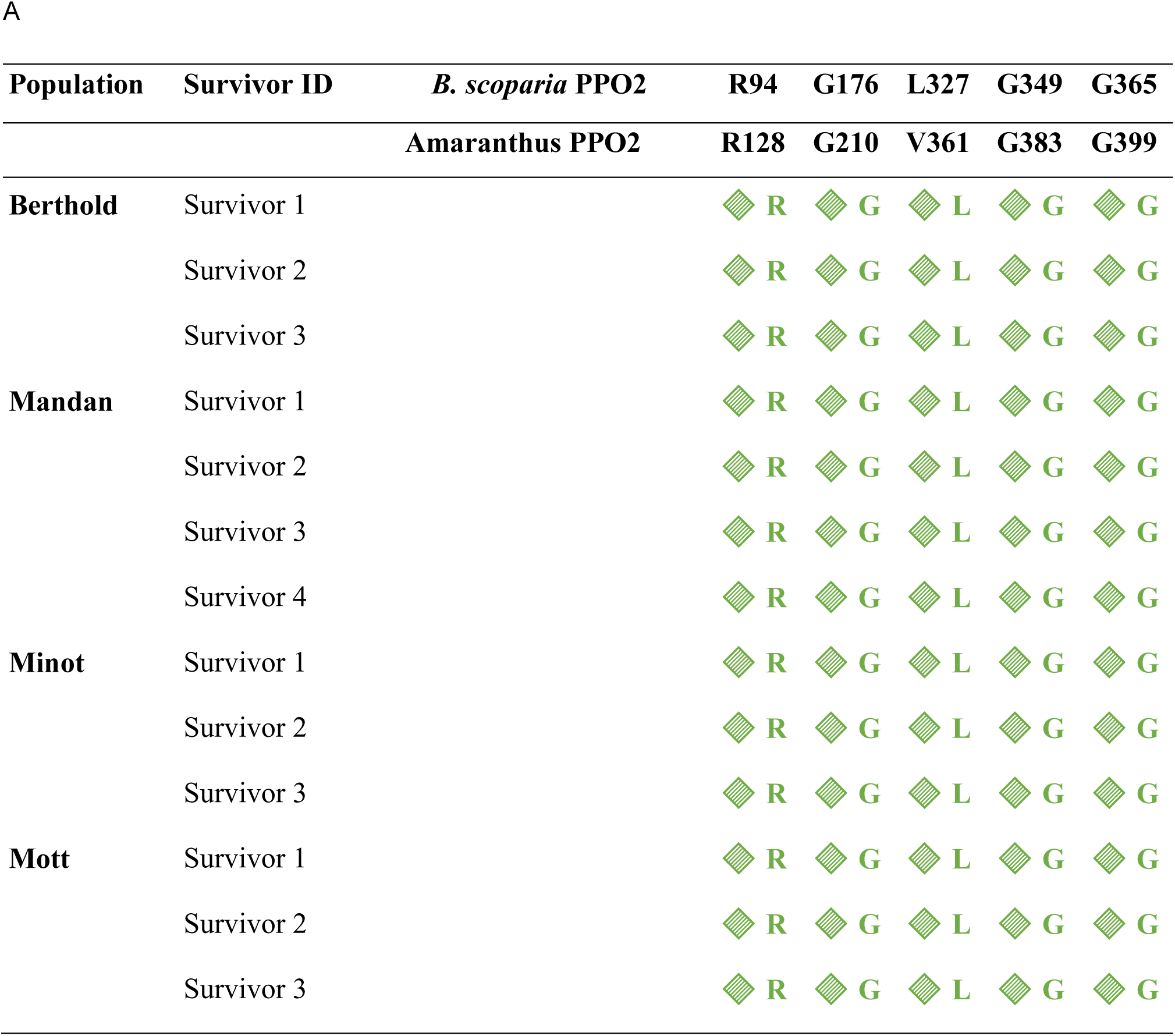
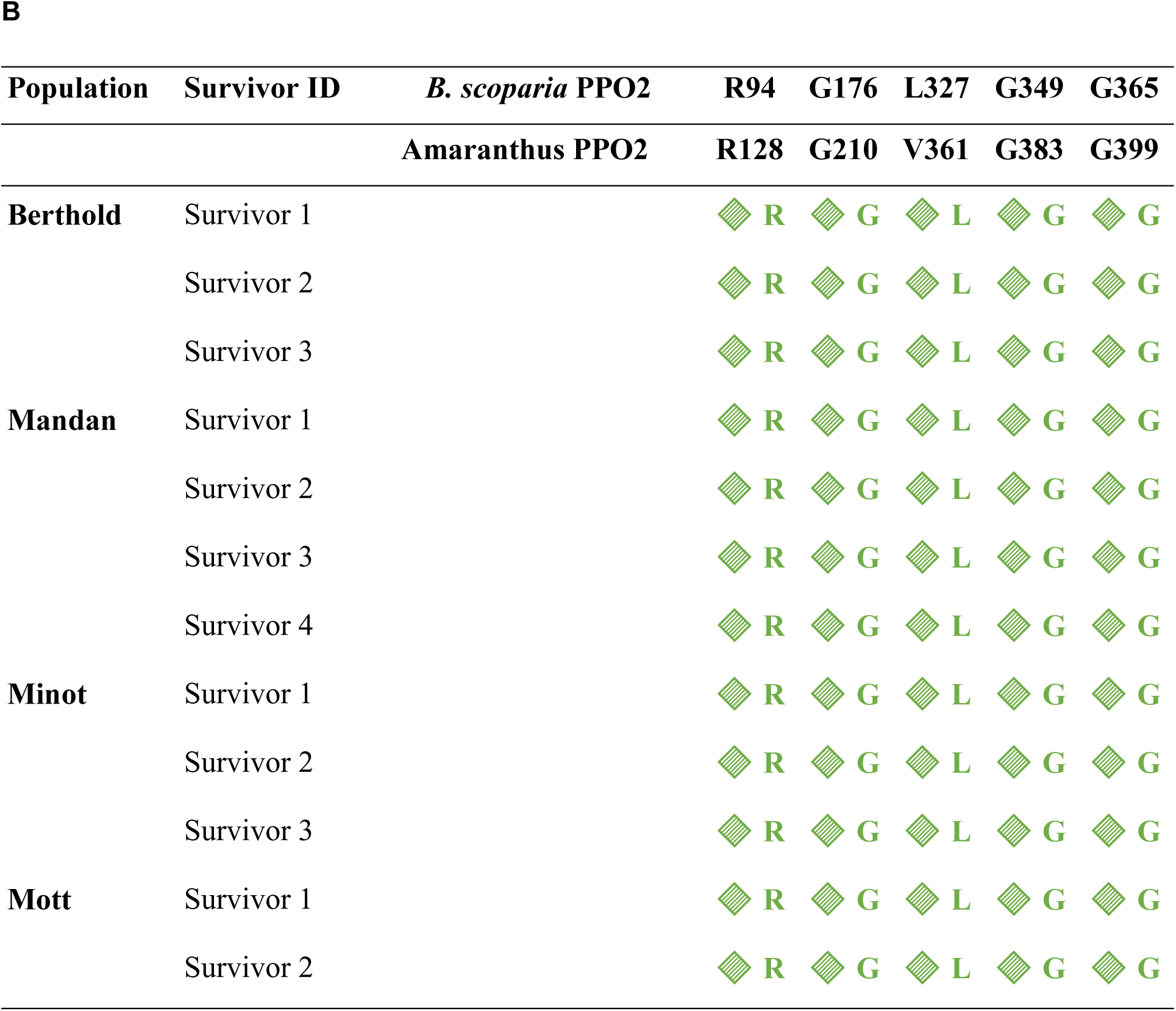
Amino acid residues at key PPO2 target-site positions in ***B. scoparia*** plants surviving treatment with PPO-inhibiting herbicides. Individual survivors from four field-collected populations (Berthold, Mandan, Minot, and Mott) were sampled after surviving labeled field rates (1X) of either (A) saflufenacil or (B) carfentrazone. Sanger sequencing was used to identify amino acid residues at known resistance-associated positions in the B. scoparia PPO2 protein (R94, G176, L327, G349, G365) and their orthologous positions in Amaranthus PPO2 (R128, G210, V361, G383, G399).

### CNV and expression analysis of *B. scoparia PPO1* and *PPO2*

TaqMan™ technology was used to determine gene copy number variation (CNV) and expression of the PPO1 and PPO2 genes in *B. scoparia* plants that survived 1X saflufenacil and 1X carfentrazone-ethyl compared with untreated susceptible plants. The TaqMan™ assays were designed to facilitate a multiplexed approach for both target and reference genes.

Digital PCR (dPCR) was performed in a final volume of 12 µL using 1,48 µL of DNA or cDNA, 0,48 µl (0.2µM) of specific primers and 0,24 µL (0.2µM) of probe (Fig S4) (biomers.net GmbH, Ulm, Germany), 3 µl of QIAcuity HighMultiplex ProbePCR Kit (Qiagen, Hilden, Germany) and 5,36 µL PCR-Grade H_2_O for the triplex dPCR. Digital PCR was performed in a dPCR thermal cycler (QIAcuity One, 5plex Device, Qiagen, Hilden Germany) in a 96-well Nanoplate with 8.500 Nanowells for each sample (QIAcuity Nanoplate 8.5k 96-well) under the following conditions: 2 min at 95°C and 55 cycles of 15 s denaturation at 95°C; 55 s annealing and elongation at 60°C. The following conditions were met during the imaging of the partitions. The FAM and HEX experiments were conducted with an exposure time of 500 milliseconds, with the gain set to 6. The ROX experiment utilized an exposure time of 400 milliseconds, also with the gain set to 6. The assessment of CNV and gene expression was conducted by employing Qiagen’s QIAcuity Software Suite, version 3.1.0.0. This process entailed the utilization of positive, negative, and no-template control wells (NTC), along with the determination of sample thresholds. Primers sequences are in table S2.

### Identification of target-site mutations in *B. scoparia* PPO1 via cDNA sequencing

To identify target-site mutations in the *B. scoparia* PPO1 gene, total RNA was extracted from young leaf tissues of plants that survived 1X saflufenacil and 1X carfentrazone-ethyl and compared with susceptible plants. Approximately 100 mg of fresh tissue was flash-frozen in liquid nitrogen and ground to a fine powder using a mortar and pestle. RNA was isolated using the RNeasy Plant Mini Kit (Qiagen, Hilden, Germany) following the manufacturer’s protocol, including an on-column DNase I treatment (Qiagen) to eliminate genomic DNA contamination. RNA integrity was assessed via 1% agarose gel electrophoresis and concentration was quantified using a NanoDrop spectrophotometer (Thermo Fisher Scientific). First-strand cDNA synthesis was performed using 1 µg of total RNA, oligo(dT) primers, and the RevertAid First Strand cDNA Synthesis Kit (Thermo Fisher Scientific) according to the manufacturer’s instructions.

The PPO1 coding sequence was amplified from cDNA using gene-specific primers designed to span the full open reading frame of *B. scoparia*. Each 25-µL PCR reaction contained 2 µL of cDNA, 0.4 µM of each primer, 200 µM dNTPs, 1× PCR buffer, 1.5 mM MgCl₂, and 1 U Taq DNA polymerase (Thermo Fisher Scientific). PCR cycling conditions were: 95°C for 3 min; 35 cycles of 95°C for 30 s, 58°C for 30 s, and 72°C for 1 min; followed by a final extension at 72°C for 7 min. Amplicons were confirmed on 1.2% agarose gels stained with SYBR Safe DNA Gel Stain (Invitrogen), and purified using the QIAquick PCR Purification Kit (Qiagen). Sequencing was performed bidirectionally using the amplification primers. Chromatograms were analyzed using Geneious Prime, and sequences were aligned to the *B. scoparia* PPO1 WT reference coding sequence. Amino acid changes resulting from nucleotide substitutions were identified to detect target-site mutations, including substitutions at position F454.

### *B. scoparia* PPO1 and PPO2 mRNA abundance during early development

The plant material of four different growth stages (germination, 2.5 cm, 6.3 cm, and 10 cm) of *B. scoparia* and *A. palmeri* (AMAPA) was homogenized using a mortar and pestle. The samples consisted of 10 plants per growth stage and were ground on dry ice. To ensure shelf life, the ground samples were stored at −80°C in Falcon tubes until RNA was extracted.

RNA was extracted from the samples using the RNeasy Plant Mini Kit I50 according to the manufacturer’s instructions. The concentration and purity of DNA were measured using a NanoDrop spectrophotometer. For cDNA synthesis, the amount of RNA extraction corresponding to 1 µg was first determined, so that 1 µg of RNA was converted to cDNA. The synthesis was performed according to the instructions of the QuantiNova Reverse Transcription Kit 200. The synthesized cDNA was stored at −20 °C until further use.

Digital PCR was performed using the dPCR system from Qiagen. Each sample was pipetted into a 96-well plate in a volume of 11.4 µL. The reaction mix was pipetted according to a template. This corresponded to a volume for 30 preparations, as 24 samples were to be measured. The reaction mixture comprised 342 µL of dPCR master mix, which contained 40 µmol µL^-1^ of specific primers, 20 µmol µL^-1^ of probes, 150.9 µL of ddH₂O and 97.5 µL of Qiagen. The master mix was then mixed with 1.6 µL DNA (1:5 diluted) sample and pipetted into the wells.

Specific fluorescent markers were used to ensure reliable detection of the *PPO1* and *PPO2* genes. In *B. scoparia*, *PPO1* was detected using fluorochrome FAM, while PPO2 was labeled HEX. In the case of *A. palmeri*, PPO1 was detected by the fluorescent dye ROX and *PPO2* by FAM. This specific labeling enabled a clear differentiation of the target genes in the respective species. Amplification was performed under the following conditions: Initial denaturation at 95°C for 2 min, followed by 40 cycles at 95°C for 10 s and 60°C for 40 s. Primers sequences are in table S2.

### Recombinant expression, purification, and *in vitro* inhibition assays of *B. scoparia* PPO1 wildtype and mutant enzymes

To investigate the functional effects of amino acid substitutions at position F454 in *B. scoparia* PPO1, three variants F454I, F454L, and F454V of the PPO1 enzyme were generated and evaluated for enzymatic activity and herbicide sensitivity. The full-length *B. scoparia* PPO1 coding sequence was synthesized *de novo* and cloned into the pRSetB expression vector (Invitrogen, Carlsbad, CA, USA) using BamHI and HindIII restriction sites. A N-terminal hexahistidine tag was included to facilitate purification. Recombinant constructs were transformed into *Escherichia coli* strain BL21(DE3)pLysS (Novagen, EMD Millipore, Billerica, MA, USA), and transformants were selected on LB agar containing 100 µg mL⁻¹ ampicillin and 34 µg mL⁻¹ chloramphenicol.

For protein expression, a single colony was used to inoculate 3 mL LB medium with antibiotics and incubated at 37°C with shaking (200 rpm) for 6 h. A 20-µL aliquot was transferred into 20 mL fresh LB medium and incubated overnight. The next day, 100 µL of the overnight culture was inoculated into 100 mL ZYM-5052 autoinduction medium (see Supplemental Data for preparation), supplemented with antibiotics, and incubated at 37°C for 5 h, followed by 21 h at 25°C.

Cells were harvested by centrifugation at 6,000 × g for 30 min at 4°C. Cell pellets were resuspended in PPO lysis buffer [10 mL g⁻¹ pellet; 50 mM NaH₂PO₄, 100 mM NaCl, 5 mM imidazole, 5% (v v^-1^) glycerol, pH 7.5, 20 mg mL⁻¹ lysozyme, 30 U mL⁻¹ DNase I] supplemented with protease inhibitors (complete EDTA-free; Roche Diagnostics, Mannheim, Germany). Suspensions were sonicated on ice (3 min, 90% amplitude, 30-s intervals). Cell debris was removed by centrifugation at 38,000 × g for 30 min at 4°C, and 2 mL of 5 M NaCl was added to the clarified supernatant.

For purification, a 500-µL bed volume of HisPur Ni-NTA resin (Thermo Fisher Scientific, IL, USA) was equilibrated with buffer (20 mM NaH₂PO₄, 50 mM NaCl, 5 mM imidazole, 5 mM MgCl₂, 17% glycerol, 0.1 mM EDTA, pH 8.0). The supernatant was passed through the resin, washed with 5.6 mL wash buffer (20 mM NaH₂PO₄, 50 mM NaCl, 5 mM imidazole, 17% glycerol, pH 7.5), and the bound protein was eluted with 1 mL elution buffer (20 mM NaH₂PO₄, 50 mM NaCl, 250 mM imidazole, 17% glycerol, pH 7.5). Protein concentrations were determined using a Scandrop nanovolume spectrophotometer (Analytikjena, Life Science, Germany). Purity and solubility were assessed by SDS-PAGE (10%) using 2.5 µg protein per lane.

PPO1 activity was assayed using a fluorescence-based method (excitation: 405 nm; emission: 630 nm) in a 187-µL reaction containing 100 mM Tris–HCl, 1 mM EDTA, 5 mM DTT, 0.0085% Tween 80, and 15 µL enzyme in resuspension buffer (50 mM Tris–HCl, pH 7.3, 3.2 mM EDTA, 20% v/v glycerol). Enzyme concentration varied by mutant to equalize maximum fluorescence in the absence of inhibitor and under saturating substrate conditions. Fluorescence units per min were calculated and normalized to WT PPO1 activity.

All PPO1 variants were subjected to dose-response assays. PPO inhibitors (Pestanal standards, Sigma-Aldrich, Steinheim, Germany) were dissolved in 80% DMSO. Ten concentrations (10 µL; range: 5 × 10⁻⁵ to 5.12 × 10⁻¹² M) were added to the reaction mixture and incubated for 30 min at room temperature. Protoporphyrinogen IX (3.24 µM; prepared from protoporphyrin via sodium amalgam reduction was then added. Fluorescence was recorded for 30.25 min (33 cycles of 55 s) using a microplate reader (CLARIOstar, BMG LabTech, Germany). Percent inhibition was calculated relative to untreated (positive) and no-enzyme (negative) controls. Assays were conducted in duplicate. IC₅₀ values (concentration of inhibitor reducing PPO1 activity by 50%) were estimated by nonlinear regression (three- or four-parameter log-logistic models). Resistance factors were calculated by dividing the IC₅₀ of each mutant variant by that of WT *B. scoparia* PPO1.

## 3. Results

### 1. Confirmation of PPO inhibitor resistance in *B. scoparia* biotypes collected from different North Dakota locations

A clear dose-dependent injury pattern was observed following saflufenacil application. The sensitive check exhibited near-complete visual injury (∼100%) across most of the treatment rates, confirming the herbicide’s effectiveness in susceptible backgrounds (Fig. 1A). At the lowest dose 12.5 g ha⁻¹ (0,5X) of saflufenacil, Berthold, Mandan, Minot, and Mott biotypes exhibited moderate injury levels below 40% while the sensitive biotype showed more than 60% injury. As expected, as the dose increased to 300 g ha⁻¹ (12X), injury progressively intensified in Berthold, Mandan, Minot, and Mott reaching approximately 60–75% for all biotypes while the sensitive plants were fully controlled, indicating lack of full herbicidal control in the four *B. scoparia* biotypes even at very high rates. At 25 g ha⁻¹ (1X) the four biotypes showed 40% injury or less, while the sensitive plants reached 90% injury, indicating herbicidal loss of efficacy at the recommended field dose rate of saflufenacil. Thus, *the B. scoparia* biotypes tested were highly resistant to saflufenacil.

**Fig. 1.**
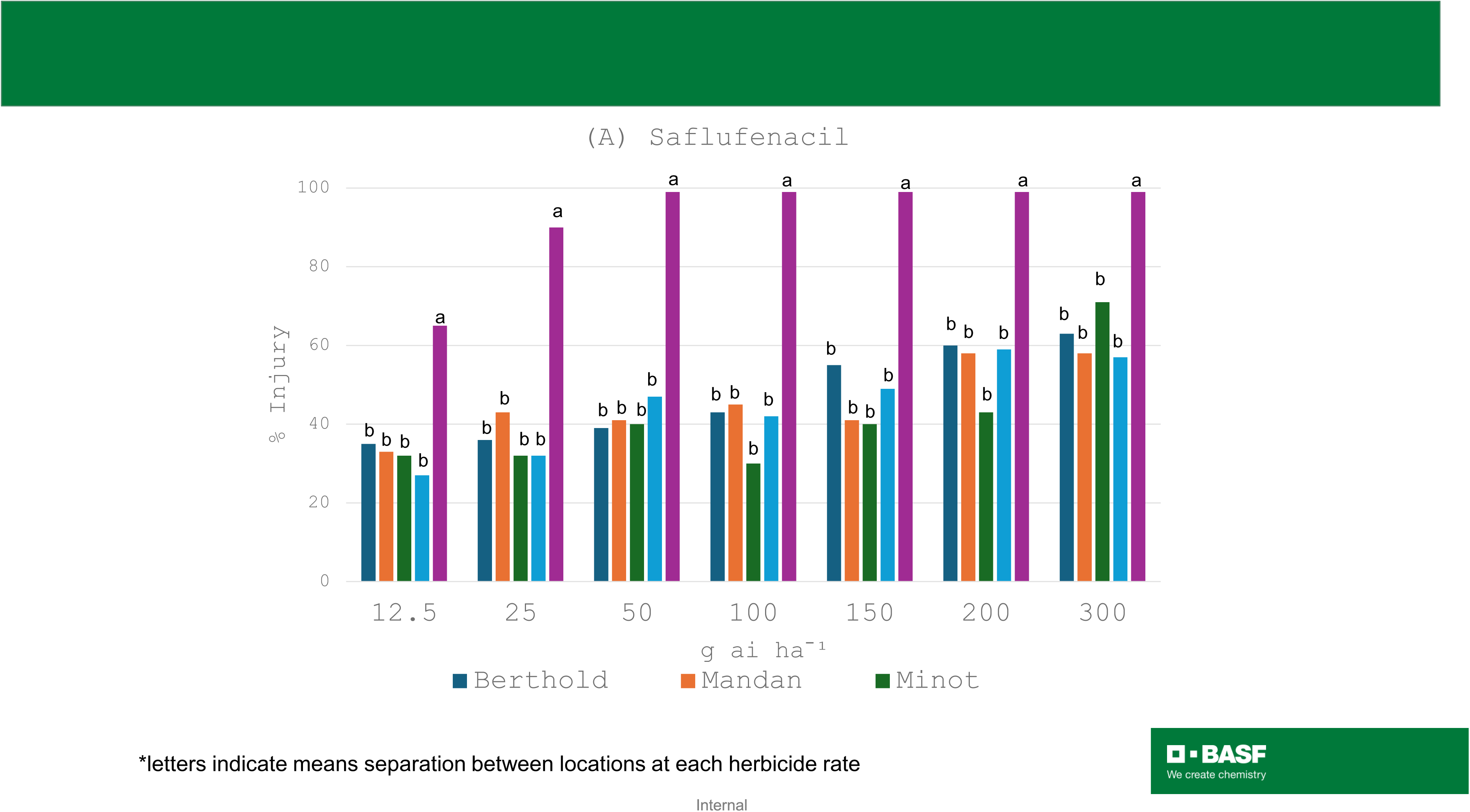
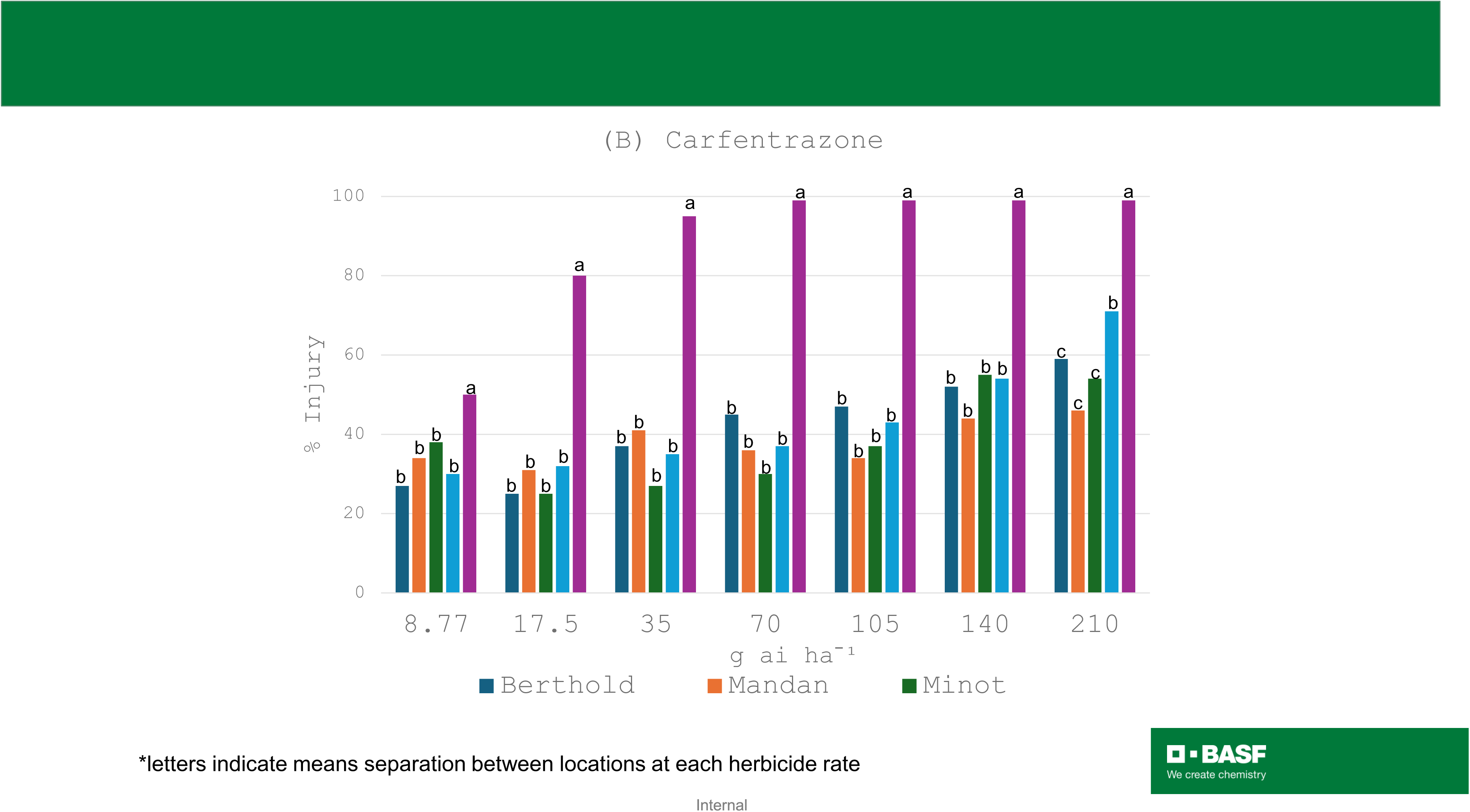
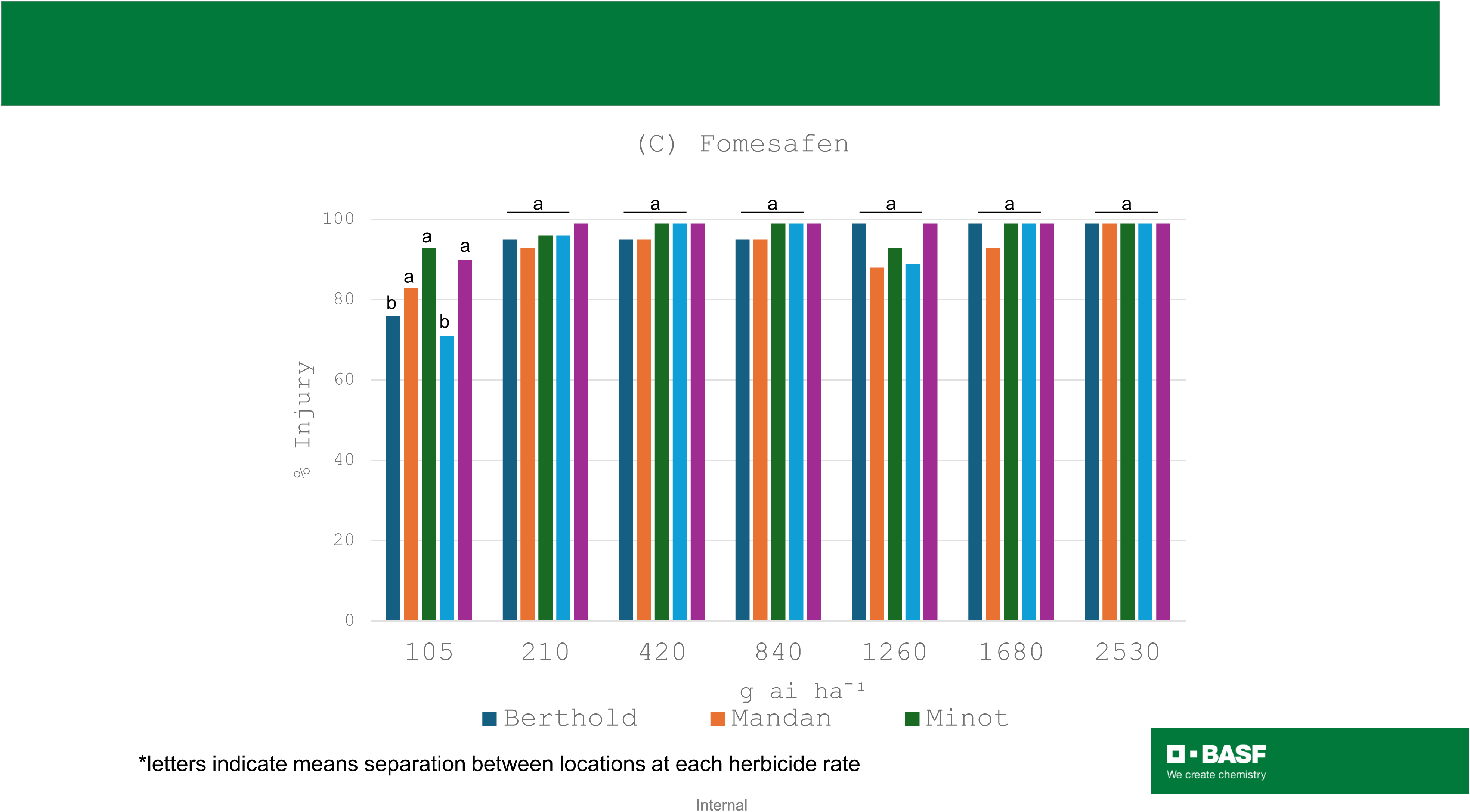
Dose–response of *B. scoparia* scoparia populations to three PPO-inhibiting herbicides under greenhouse conditions. (A) saflufenacil, (B) carfentrazone, and (C) fomesafen were applied at increasing rates (x-axis, g ai ha⁻¹) to four field-collected populations (Berthold, Mandan, Minot, and Mott) and a sensitive reference population was used as control. The lowest herbicide rate (leftmost value on the x-axis) corresponds to 0.5X the labeled field use rate. Visual injury was assessed 14 days after treatment. Different letters indicate statistically significant differences among populations at each herbicide rate (one-way ANOVA followed by post hoc mean separation, α = 0.05).

Responses to carfentrazone-ethyl were similar to those observed for saflufenacil. The sensitive check once again showed severe injury (∼100%) across most rates, confirming herbicidal activity (Fig. 1B). At the lowest dose of 8.77 g ha⁻¹, injury among the biotypes ranged between 25–40%. With increasing doses, injury levels rose in all biotypes and at the recommended field dose rate of 17.5 g ha⁻¹ (1X) all four B. scoparia biotypes showed injury less than 40%, while the sensitive plants reached 80% injury, demonstrating significant resistance to carfentrazone-ethyl.

Fomesafen treatments led to high levels of visual injury across all biotypes, with little differentiation among them (Fig. 1C). Even at the lowest dose of 105 g ha⁻¹ (0.5X), injury levels were significant for all tested biotypes, closely matching the response of the sensitive check. As the dose increased to 2,530 g ha⁻¹, the injury remained uniformly high, with all biotypes approaching full damage (∼95–100%). At the 210 g ha⁻¹ (1X) field rate, all biotypes and the sensitive standard were near-fully controlled with no statistical difference among them. This uniformity in response suggests a general susceptibility to fomesafen in the tested biotypes, with minimal variability. These data together strongly indicate that the tested *B. scoparia* biotypes were highly resistant to carfentrazone-ethyl and saflufenacil, while fomesafen retained high herbicidal efficacy.

### 2. Copy number and expression of PPO1 and PPO2 remains unchanged in resistant *B. scoparia* plants

To assess the potential involvement of gene amplification in PPO inhibitor resistance, CNV of PPO1 and PPO2 was evaluated in *B. scoparia* plants that survived 1X treatment of saflufenacil and carfentrazone-ethyl relative to a sensitive control. As shown in Figure 2A, no consistent increase in copy number was observed for both genes across resistant individuals from the four biotypes. Fold-change values clustered around or below unity, indicating that PPO1 and PPO2 were not genomically amplified in the resistant plants. Thus, CNV is unlikely to be a major mechanism contributing to PPO inhibitor resistance in these *B. scoparia* plants. In addition, quantification of PPO1 and PPO2 transcript levels across *B. scoparia* survivor plants revealed limited variation in gene expression with most samples showing mRNA abundance levels comparable to the susceptible reference, with only minor fluctuations observed between treatments or locations (Fig. 2B). Together, these results suggest that PPO1 and PPO2 expression remain stable in *B. scoparia* survivor plants indicating that neither CNV nor expression changes in the PPO1 and PPO2 isoforms were likely to mediate resistance to saflufenacil and carfentrazone-ethyl.

**Fig. 2.**
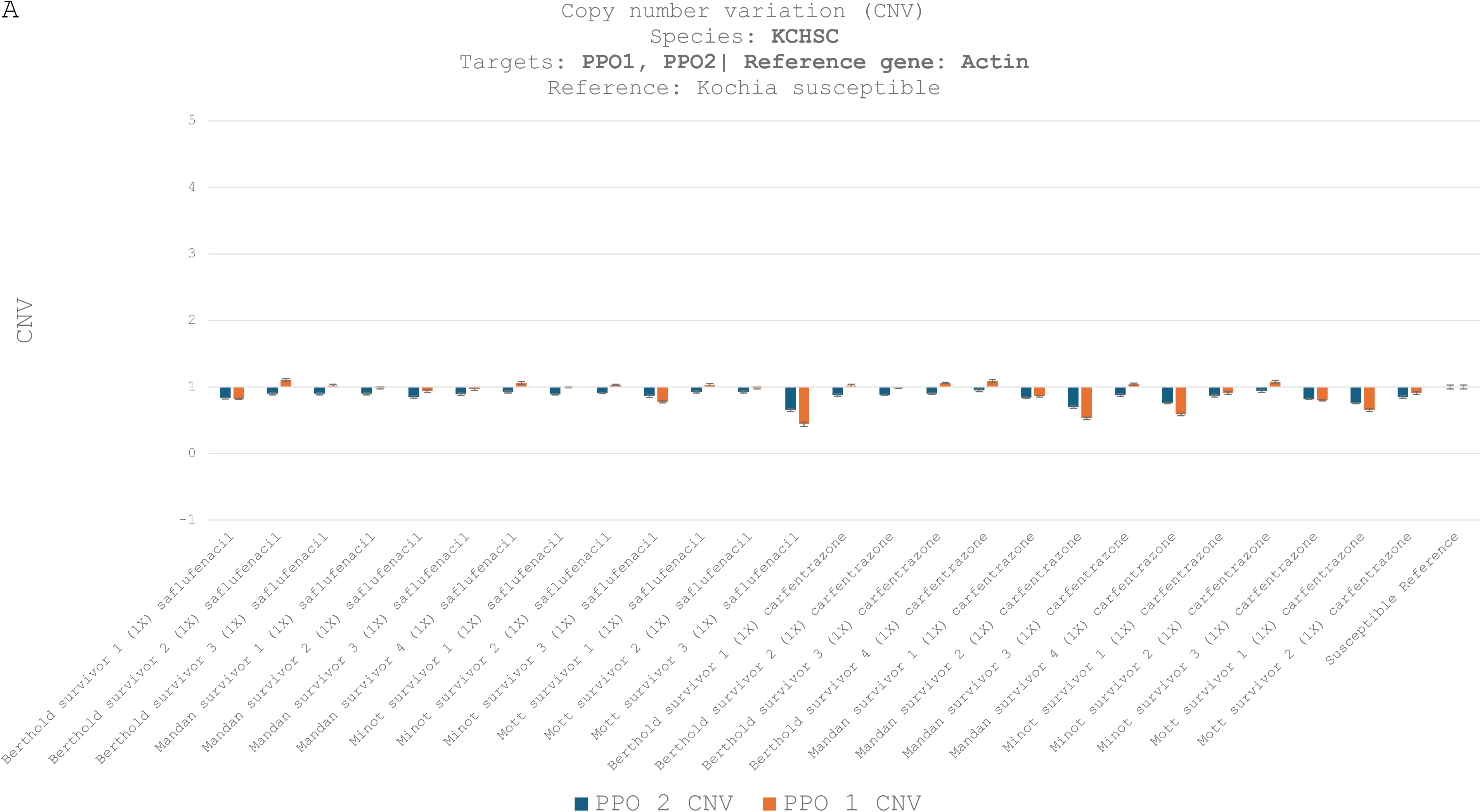
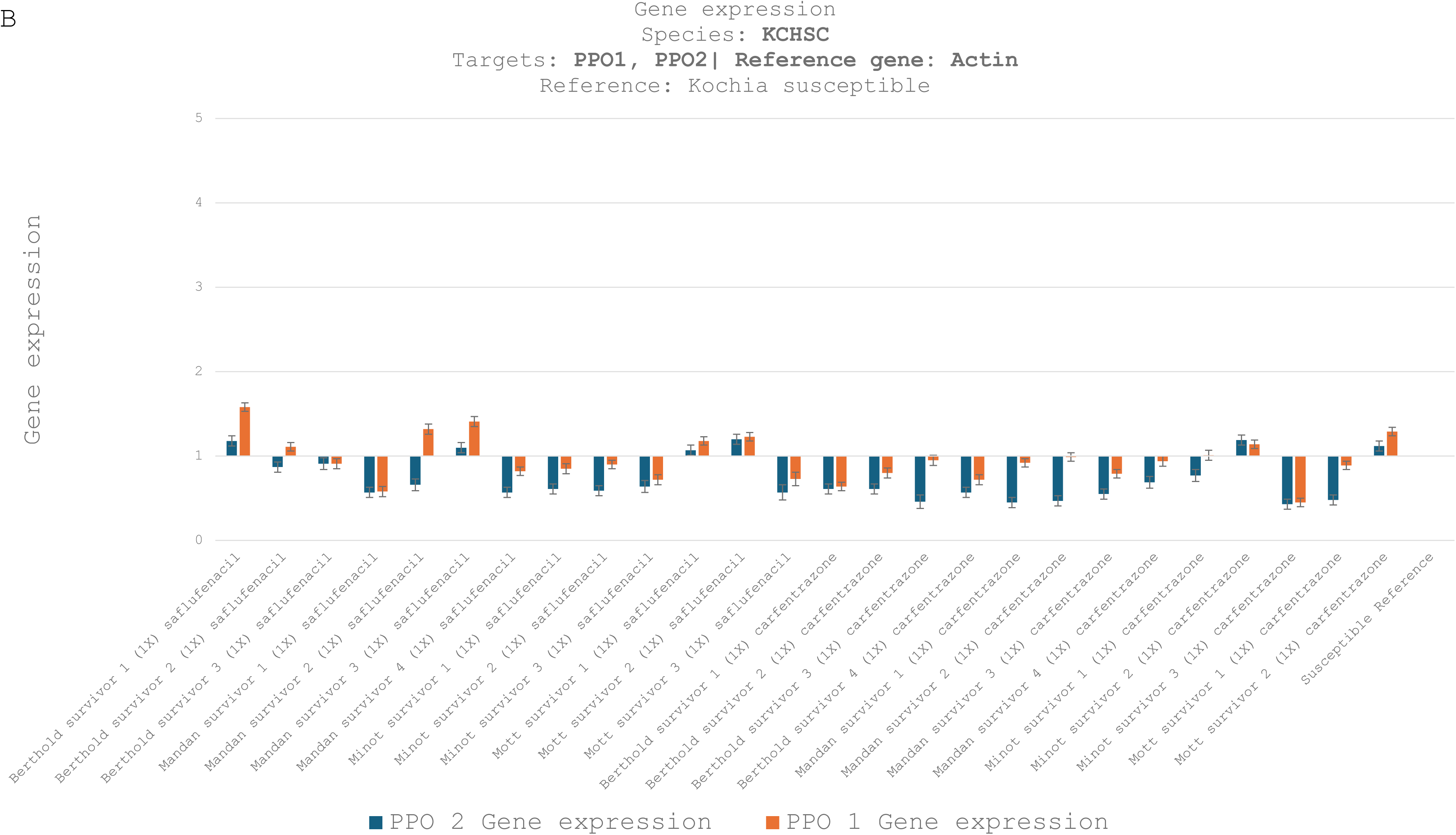
Copy number variation (CNV) and gene expression of PPO1 and PPO2 in *B. scoparia* individuals surviving treatment with PPO-inhibiting herbicides. (A) CNV was assessed by digital qPCR using actin as a reference gene, and values were normalized to a sensitive reference population (set to 1.0). (B) Gene expression was measured by digital qPCR in the same individuals, normalized to actin and relative to the sensitive population. Plants were collected following survival to labeled field rates (1X) of either saflufenacil or carfentrazone. Bars represent 95% confidence intervals (CI95)

### 3. PPO1 F454 mutations are associated with resistance in *B. scoparia* survivor plants

To elucidate the molecular basis of resistance, PPO1 and PPO2 cDNAs were sequenced in survivor plants following treatment with 1X rates of saflufenacil or carfentrazone-ethyl. Sanger sequencing was employed to identify possible amino acid substitutions derived from nucleotide changes.

Sequencing of the PPO2 gene revealed complete conservation of amino acid residues at all known target-site positions (R94, G176, L327, G349, and G365 in *B. scoparia* PPO2; R128, G210, V361, G383, and G399 in *Amaranthus* PPO2 homologs). As shown in Tables 1A and 1B, all survivor plants retained WT residues across these positions, regardless of herbicide treatment or biotype origin. This absence of non-synonymous substitutions indicates that PPO2 variations in these key target site positions did not occur in the survivor plants. Substitutions at PPO2 positions I201 and V331 were detected in *B. scoparia* survivor plants treated with 1X saflufenacil Supp. Table 1 (A) or carfentrazone-ethyl Supp. Table 1 (B). While I201 remained WT in most individuals, I/L heterozygosity was observed in two Mandan and one Berthold survivors, and a homozygous L201 variant was exclusive to two Mott survivors. At V331, most plants retained the WT valine. However, V/F heterozygotes were found in one Minot and all Mott survivors. The inconsistent occurrence of such amino acid substitutions indicates that they were unlikely to contribute to the resistant phenotype of the survivor plants. No further amino acid substitutions were detected in PPO2.

In contrast, sanger sequence analysis of the PPO1 gene revealed a diverse array of amino acid substitutions consistently at position F454. As presented in Table 2A and B, all survivor plants from both treatment groups exhibited either homozygous or heterozygous mutations at this site, including F/I, F/L, I/L, I, L, and V substitutions. The F454I mutation emerged as the most prevalent across populations, particularly in individuals from Berthold and Mandan, where it was frequently detected in a heterozygous (F/I) or homozygous (I) state; especially all three Berthold survivors and all four Mandan individuals surviving carfentrazone-ethyl.

**Table 2.**
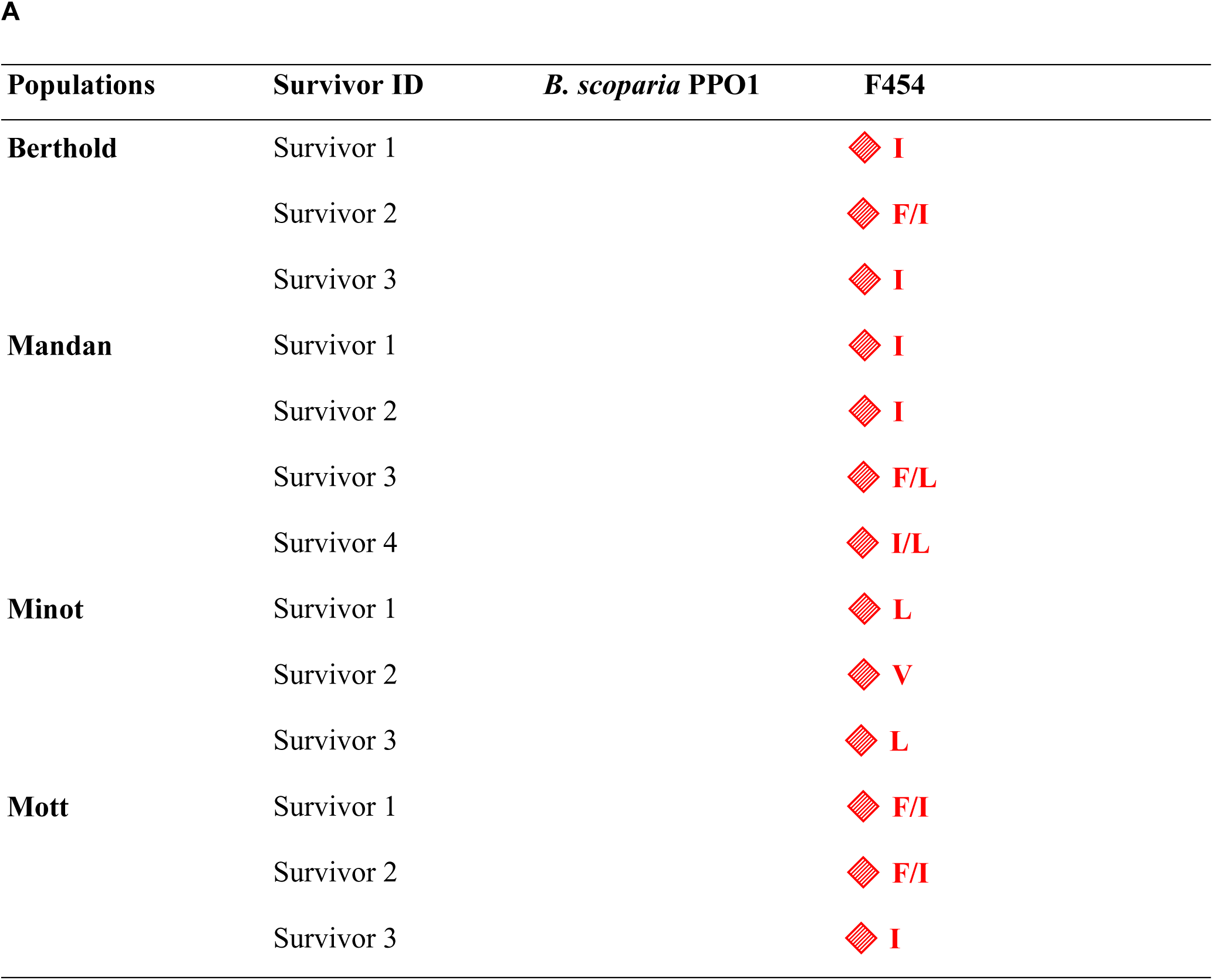
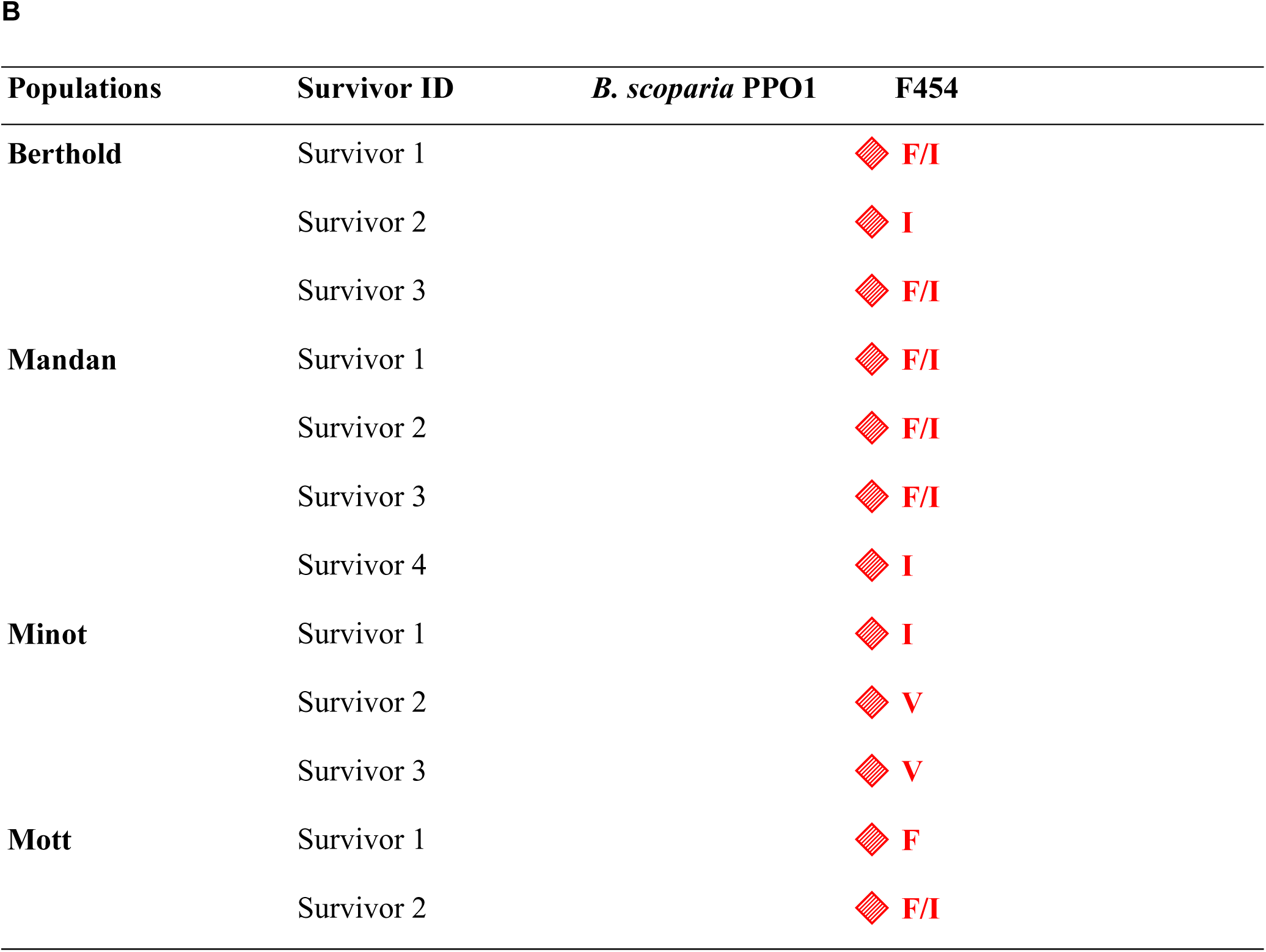
Amino acid residues at the F454 position of PPO1 in ***B. scoparia*** survivors to PPO-inhibiting herbicide treatment. Individual plants from four field-collected populations (Berthold, Mandan, Minot, and Mott) were sampled after surviving 1X labeled field rates of either (A) saflufenacil or (B) carfentrazone. Sanger sequencing was used to identify amino acid substitutions at position F454 of the B. scoparia PPO1 protein. Residues are shown for each survivor. Where two amino acids are listed (e.g., F/I), this indicates heterozygosity at the locus, with one allele carrying the wild-type residue and the other the substitution.

**A**
| Populations | Survivor ID | *B. scoparia* PPO1 | F454 |
| --- | --- | --- | --- |
| **Berthold** | Survivor 1 |  | ◆ **I** |
|  | Survivor 2 |  | ◆ **F/I** |
|  | Survivor 3 |  | ◆ **I** |
| **Mandan** | Survivor 1 |  | ◆ **I** |
|  | Survivor 2 |  | ◆ **I** |
|  | Survivor 3 |  | ◆ **F/L** |
|  | Survivor 4 |  | ◆ **I/L** |
| **Minot** | Survivor 1 |  | ◆ **L** |
|  | Survivor 2 |  | ◆ **V** |
|  | Survivor 3 |  | ◆ **L** |
| **Mott** | Survivor 1 |  | ◆ **F/I** |
|  | Survivor 2 |  | ◆ **F/I** |
|  | Survivor 3 |  | ◆ **I** |

**B**
| **Populations** | **Survivor ID** | ***B. scoparia* PPO1** | **F454** |
| --- | --- | --- | --- |
| **Berthold** | Survivor 1 |  | ◆ **F/I** |
|  | Survivor 2 |  | ◆ **I** |
|  | Survivor 3 |  | ◆ **F/I** |
| **Mandan** | Survivor 1 |  | ◆ **F/I** |
|  | Survivor 2 |  | ◆ **F/I** |
|  | Survivor 3 |  | ◆ **F/I** |
|  | Survivor 4 |  | ◆ **I** |
| **Minot** | Survivor 1 |  | ◆ **I** |
|  | Survivor 2 |  | ◆ **V** |
|  | Survivor 3 |  | ◆ **V** |
| **Mott** | Survivor 1 |  | ◆ **F** |
|  | Survivor 2 |  | ◆ **F/I** |

**Table 3.** IC_50_ values of *B. scoparia* PPO1 (WT), F454V, L and I enzymes. IC_50_ is expressed in molar concentrations which represents the amount of a given herbicide that inhibits 50% of the recombinant protein activity *in vitro.* % Inhibition at 10^-4^M represents the total enzyme inhibition with the highest herbicide concentration used in the assay. Resistance factor (RF) was calculated as IC₅₀(mutant) / IC₅₀(WT).

|  | IC<sub>50</sub> [M] | % Inhibition at 10<sup>-4</sup>M | RF |
| --- | --- | --- | --- |
| Saflufenacil |  |  |  |
| KSHCS PPO1 WT | 8.60E-07 | 90 | - |
| KSHCS PPO1 F454V | 1.97E-05 | 73 | 23 |
| KSHCS PPO1 F454L | 4.61E-06 | 82 | 5 |
| KSHCS PPO1 F454I | 4.96E-06 | 80 | 6 |
| Carfentrazone |  |  |  |
| KSHCS PPO1 WT | 5.20E-07 | 88 | - |
| KSHCS PPO1 F454V | 2.88E-05 | 53 | 55 |
| KSHCS PPO1 F454L | 1.17E-05 | 55 | 23 |
| KSHCS PPO1 F454I | 1.34E-05 | 52 | 26 |
| Fomesafen |  |  |  |
| KSHCS PPO1 WT | 8.06E-08 | 98 | - |
| KSHCS PPO1 F454V | 3.49E-08 | 95 | 0.4 |
| KSHCS PPO1 F454L | 1.22E-08 | 98 | 0.2 |
| KSHCS PPO1 F454I | 3.67E-08 | 99 | 0.5 |

Evidence of allelic diversity was observed not only among populations but also within them. For instance, Minot survivors exposed to saflufenacil displayed genotypic heterogeneity at F454, including individuals carrying V, L substitutions. The presence of multiple amino acid variants at a single residue suggests a high degree of selective pressure on this locus and points to the evolution of distinct mutational pathways associated with resistance. No other amino acid substitutions were detected in the PPO1 isoform.

Taken together, these results suggest that substitutions at position F454 in PPO1, F454I, F454L, and F454V are associated with the resistance phenotype in the four *B. scoparia* biotypes. In contrast, no evidence supports a role for PPO2 in herbicide resistance in these populations. The observed prevalence, allelic diversity, and zygosity of F454 substitutions across survivors from both saflufenacil and carfentrazone-ethyl further support the critical involvement of these substitutions in conferring resistance.

### 4. F454 substitutions reduce PPO1 sensitivity to saflufenacil and carfentrazone but not fomesafen *in vitro*

To elucidate the impact of amino acid substitutions at position F454 on the sensitivity of *B. scoparia* PPO1 to PPO-inhibiting herbicides, we evaluated the inhibitory responses of recombinant PPO1 WT and mutant enzymes (F454V, F454L, and F454I) to saflufenacil, carfentrazone-ethyl, and fomesafen. IC₅₀ values were determined as a measure of half-maximal inhibition, and percent inhibition was assessed at a saturating herbicide concentration (10⁻⁴ M).

For saflufenacil, the WT enzyme displayed an IC₅₀ of 8.60 × 10⁻⁷ M, with 90% inhibition at 10⁻⁴ M. Substitution with valine at position F454 (F454V) markedly reduced herbicide sensitivity, increasing the IC₅₀ to 1.97 × 10⁻⁵ M and yielding a resistance factor (RF) of 23. In contrast, the F454L and F454I variants exhibited moderate resistance, with IC₅₀ values of 4.61 × 10⁻⁶ M and 4.96 × 10⁻⁶ M, corresponding to RFs of 5 and 6, respectively. At 10⁻⁴ M of saflufenacil both mutants showed inhibition ranging 80–82%, indicating lack of full inhibition under high herbicide concentrations. Resistance was even more pronounced in response to carfentrazone-ethyl. The WT enzyme showed an IC₅₀ of 5.20 × 10⁻⁷ M, with 88% inhibition at 10⁻⁴ M. However, the F454V variant exhibited a 55-fold increase in resistance (IC₅₀ = 2.88 × 10⁻⁵ M), while F454L and F454I mutants showed IC₅₀ values of 1.17 × 10⁻⁵ M and 1.34 × 10⁻⁵ M, respectively, with corresponding RFs of 23 and 26. Consistently, inhibition at 10⁻⁴ M was substantially diminished across all three mutants (55–26%), underscoring a substantial loss of carfentrazone-ethyl efficacy in the presence of F454 substitutions. By contrast, fomesafen retained high inhibitory potency across all enzyme variants. The WT PPO1 enzyme exhibited an IC₅₀ of 8.06 × 10⁻⁸ M and 98% inhibition at 10⁻⁴ M. Notably, the F454V, F454L, and F454I substitutions had minimal impact, with IC₅₀ values ranging from 1.22 × 10⁻⁸ M to 3.67 × 10⁻⁸ M and RFs between 0.2 and 0.5. Inhibition at 10⁻⁴ M remained consistently high (95–99%) across all enzyme variants, indicating that fomesafen efficacy is largely unaffected by these target-site substitutions.

The above experiments suggest herbicide-specific effects of F454 mutations on PPO1 inhibition. While F454 substitutions substantially impair enzyme sensitivity to saflufenacil and carfentrazone-ethyl, they exert negligible influence on fomesafen activity, highlighting distinct structural interactions between PPO1 substitutions and these herbicidal chemistries.

### 5. Ectopic expression of F454L, I and V in *Arabidopsis* transgenic lines

To analyze whether amino acid substitutions at position F454 in PPO1 are sufficient to confer tolerance to saflufenacil, fomesafen or carfentrazone-ethyl *in planta*, we ectopically expressed the *B. scoparia PPO1* as WT protein or with F454 substitutions to leucin, isoleucine or valine in the model organism *Arabidopsis thaliana* and assessed the sensitivity of the transgenics to PPO inhibitors. Transgenic lines were treated with the 2X, 1X and 0.5X field dose rates, respectively. Three to five independent T1 transgenic lines expressing the different *PPO1* variants were tested. Based on the visual assessment, ectopic expression of WT *B. scoparia PPO1* did not increase tolerance to saflufenacil and fomesafen at all tested rates. (Figure 3 and Figure S1). For carfentrazone-ethyl, WT *B. scoparia PPO1* expression led to a slight reduction in sensitivity at 0.5X field rate which was observed in only one transgenic line but did not confer tolerance at higher rates (Figure 3 and Figure S1). In contrast, for saflufenacil and carfentrazone-ethyl two or more independent lines expressing PPO1 F454I, F454L or F454V showed a dramatic increase of tolerance. Notably, saflufenacil tolerance was observed for F454I and F454V substitutions at all rates, whereas no tolerant transgenic lines were observed for the F454L substitution at the 2X rates. For carfentrazone-ethyl, lines expressing PPO1 F454I, F454L or F454V also conferred tolerance at all rates tested (Figure 3 and Figure S1). Conversely, at all tested rates for fomesafen, expressions of the PPO1 F454 mutant variants in transgenic lines did not confer tolerance to fomesafen at all rates tested (Figure 3 and Figure S1). Taken together, these findings support that PPO1 variants with F454 substitutions endow resistance to PPO-inhibiting herbicides saflufenacil and carfentrazone-ethyl, while fomesafen efficacy was unaffected.

**Fig. 3.**
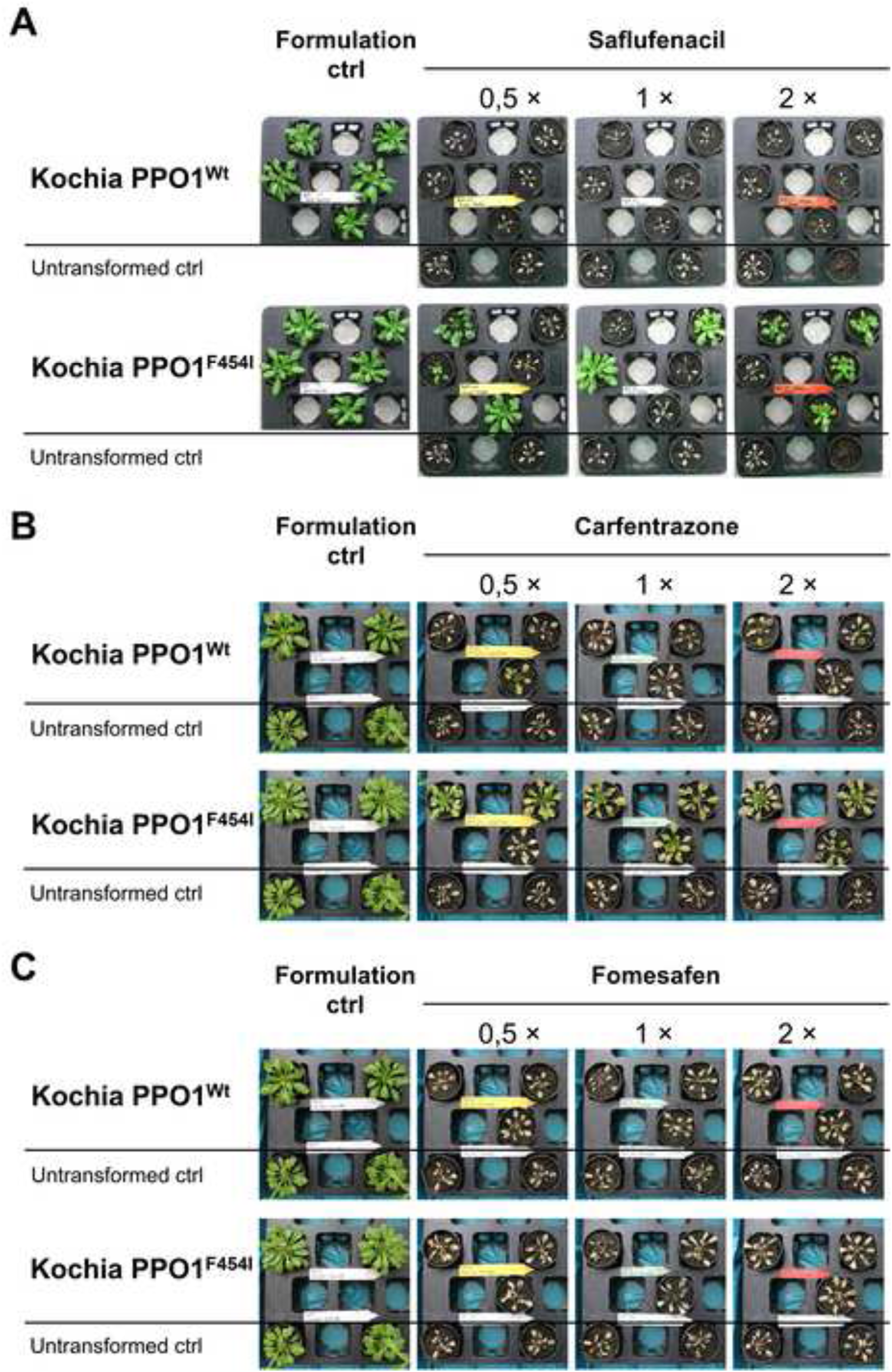
*Arabidopsis* lines in the T1 Generation ectopically expressing indicated *B. scoparia PPO1* variants were sprayed with saflufenacil, carfentrazone or fomesafen. The upper three to five pots contain independent transgenic events (T1 plants, were selected with imazamox to confirm the presence of resistance gene T-DNA via AHAS herbicide tolerance) the two plants in the lowest row represent the untransformed MC24 wild type control. (A) Images are shown of *Arabidopsis* lines ectopically expressing *B. scoparia PPO^Wt^*, *B. scoparia PPO^F454I^* (upper three rows of trays) or the untransformed control (lowst row of trays) after spray treatment with the formulation control or saflufenacil (0.5 ×: 25 g a.i./ha, 1 ×: 50 a.i./ha or 2 ×: 100 a.i./ha field use rates) 11 days after treatment. (B) Images are shown of *Arabidopsis* lines ectopically expressing *B. scoparia PPO^Wt^*, *B. scoparia PPO^F454I^* (upper two rows of trays) or the untransformed control (lowst row of trays) after spray treatment with the formulation control or carfentrazon (0.5 ×: 210 g a.i./ha, 1 ×: 420 a.i./ha or 2 ×: 820 a.i./ha field use rates) 15 days after treatment. (C) Images are shown of *Arabidopsis* lines ectopically expressing *B. scoparia PPO^Wt^*, *B. scoparia PPO^F454I^* (upper two rows of trays) or the untransformed control (lowst row of trays) after spray treatment with the formulation control or fomesafen (0.5 ×: 17.5 g a.i./ha, 1 ×: 35 a.i./ha or 2 ×: 70 a.i./ha field use rates) 15 days after treatment.

### 6. Molecular modeling of PPO-inhibiting herbicides in *B. scoparia* PPO1 mutant enzymes

Fomesafen, carfentrazone-ethyl and saflufenacil exhibited different PPO1 inhibitory efficacy in the presence of target-site substitutions at position F454, with fomesafen retaining activity against the F454V, F454L, and F454I variants, while carfentrazone-ethy; and saflufenacil lose inhibitory potency.

To understand the molecular basis of this differential efficacy, we constructed a homology model of *B. scoparia* PPO1 based on an inhouse co-crystal structure of *Eleusine indica* PPO1 with an inhibitor. The *B. scoparia* PPO1 showed 69% sequence identity and 82% similarity (BLOSUM62) with the *Eleusine* template. Notably, the only residue in direct contact with the herbicide binding site that differs between the two species is phenylalanine (F454 in *B. scoparia*) corresponding to tyrosine (Y418) in *Eleusine*. This model enabled a comparative docking analysis of herbicide binding to both WT and mutant PPO1 variants.

Docking simulations were performed using the *B. scoparia* PPO1 homology model, incorporating structural data from multiple PPO inhibitor complexes, including *Nicotiana tabacum* PPO2 (PDB: 1SEZ), ELEIN PPO1, and PPO2 structures co-crystallized with saflufenacil from *A. tuberculatus*. All three herbicides’ binding modes align with typical PPO inhibitor characteristics, with their aromatic warheads—trifluoromethylphenyl (fomesafen), uracil (saflufenacil), and triazole (carfentrazone)—positioned between helix P241–A246 and residue F454, forming canonical Π–Π interactions (Fig. 4).

**Fig. 4.**
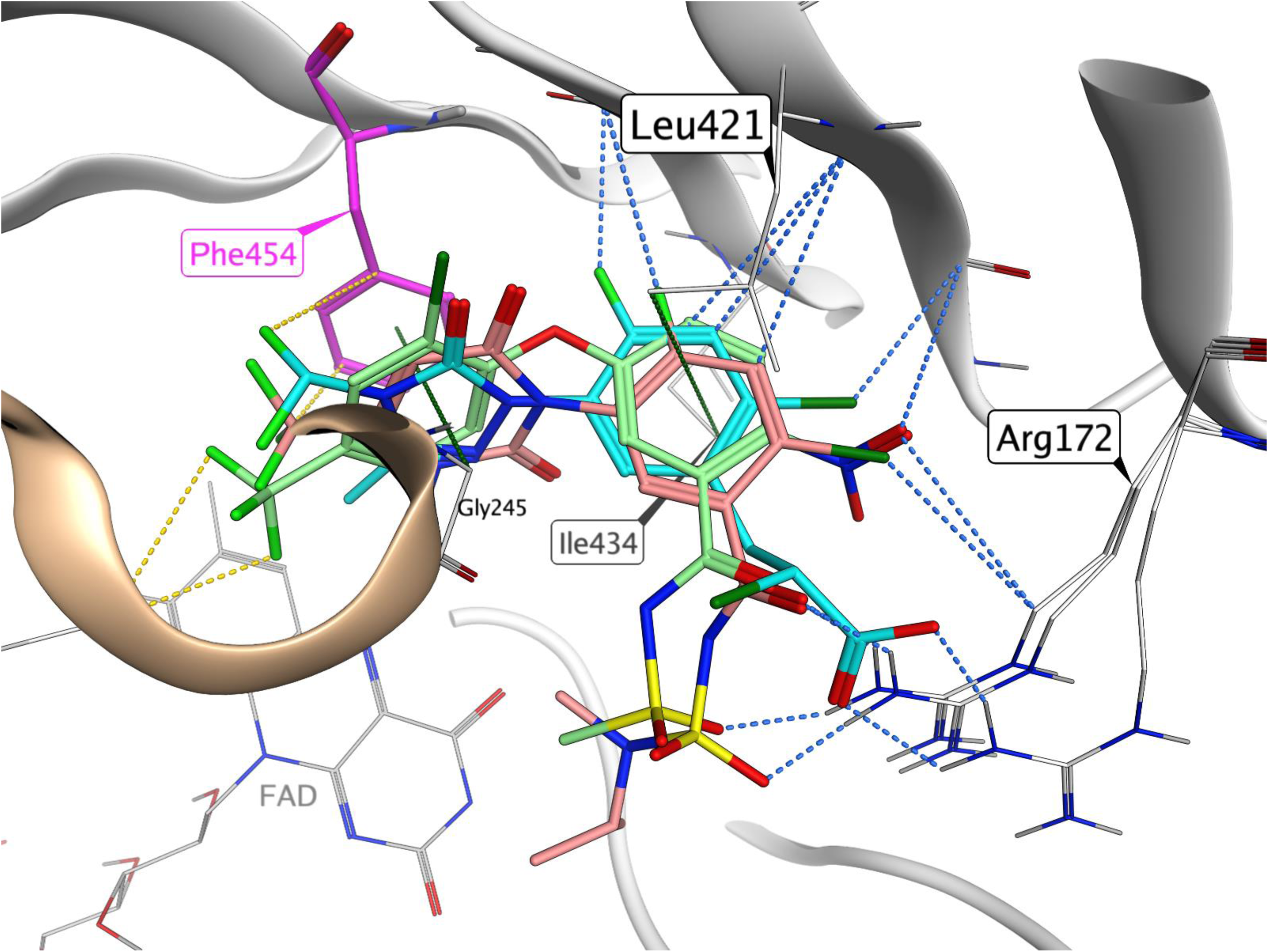
Common Ligand Interactions of PPO Herbicides in the *B. scoparia* PPO1 Homology Model. The secondary structure of the protein is illustrated in a white cartoon format. The FAD cofactor is represented with grey lines. Oxygen atoms are marked in red, nitrogen atoms in blue, sulfur atoms in yellow, chlorine atoms in dark green, and fluorine atoms in light green. Within the binding pocket, the three herbicides are displayed in stick form: Fomesafen (green carbon atoms), Carfentrazone (cyan carbon atoms), and Saflufenacil (red carbon atoms) are overlaid. The warheads are positioned between Phe454 (magenta carbon atoms) and Gly245 from the alpha helix (gold), allowing for interactions with these amino acids through pi-pi and sigma-pi interactions (green dashed lines). The fluorine group substituents can engage in favorable van der Waals interactions (yellow dashed lines) in this section of the pocket. The scaffolds are situated between Ile434 and Leu421 and are stabilized by sigma-pi interactions (green dashed lines). Additionally, the scaffolds and side chains can interact with the beta sheets of PPO1 through multipolar interactions or charge-assisted hydrogen bonds (blue dashed lines) with Arg172. Arg172 is depicted in different orientations due to its flexible side chain, which can adjust to accommodate the respective herbicide.

Fomesafen binds to F454 at an angle about 18 degrees different from carfentrazone-ethyl and saflufenacil, which does not impact the Π–Π interactions, as F454 can adjust to this variation. However, it results in different exit paths for the CF3 groups (and CF2 for carfentrazone-ethyl), which ileads to notable hydrophobic interactions to the binding pocket. These interactions to F454 are more pronounced for carfentrazone-ethyl and saflufenacil.

Across all herbicides, the molecular scaffolds were twisted by ∼60° relative to their warheads, positioning the second ring systems between residues L421 and I434, supporting σ–Π interactions (Fig. 4). The scaffolds can also form multipolar interactions with halogens or polarized hydrogen bonds towards beta-sheet secondary structures. In particular, the nitro group of fomesafen fits well in the binding pocket near the side chain of R172. Furthermore, fomesafen diphenyl ether scaffold includes an additional rotatable bond, providing increased conformational flexibility compared to the rigid heterocyclic frameworks of carfentrazone-ethyl and saflufenacil. The side chains are exposed to the solvent and can interact with R172 through charge-supported hydrogen bonds. Comparing the three herbicides, differences can be identified between the binding mode of fomesafen to carfentrazone-ethyl and saflufenacil in PPO1. The F454V, F454L, and F454I substitutions have profound impacts on herbicide binding, primarily by disrupting two critical interaction aspects. First, these mutations abolish Π–Π stacking interactions that are especially important for carfentrazone-ethyl and saflufenacil, both of which possess electron-deficient warheads well-suited to such interactions (Fig. 5). While fomesafen also favors aromatic contacts, its electron-rich phenyl ring allows for σ–Π interactions with aliphatic side chains introduced by the mutations. Second, the substitutions introduce steric alterations to the binding pocket. The branched, hydrophobic side chains of valine, leucine, and isoleucine reshape the warhead binding site, affecting additionally the placement of the scaffold. This spatial rearrangement weakens interactions between the scaffold and the β-sheet backbone, which are important for stabilizing inhibitor binding. Fomesafen, with its more flexible ether linkage, can better accommodate these changes by adopting alternative conformations keeping favorable shape complementarity within the altered pocket. In contrast, the rigidity of carfentrazone-ethyl and saflufenacil limits their ability to adapt to the new binding site geometry.

**Fig. 5.**
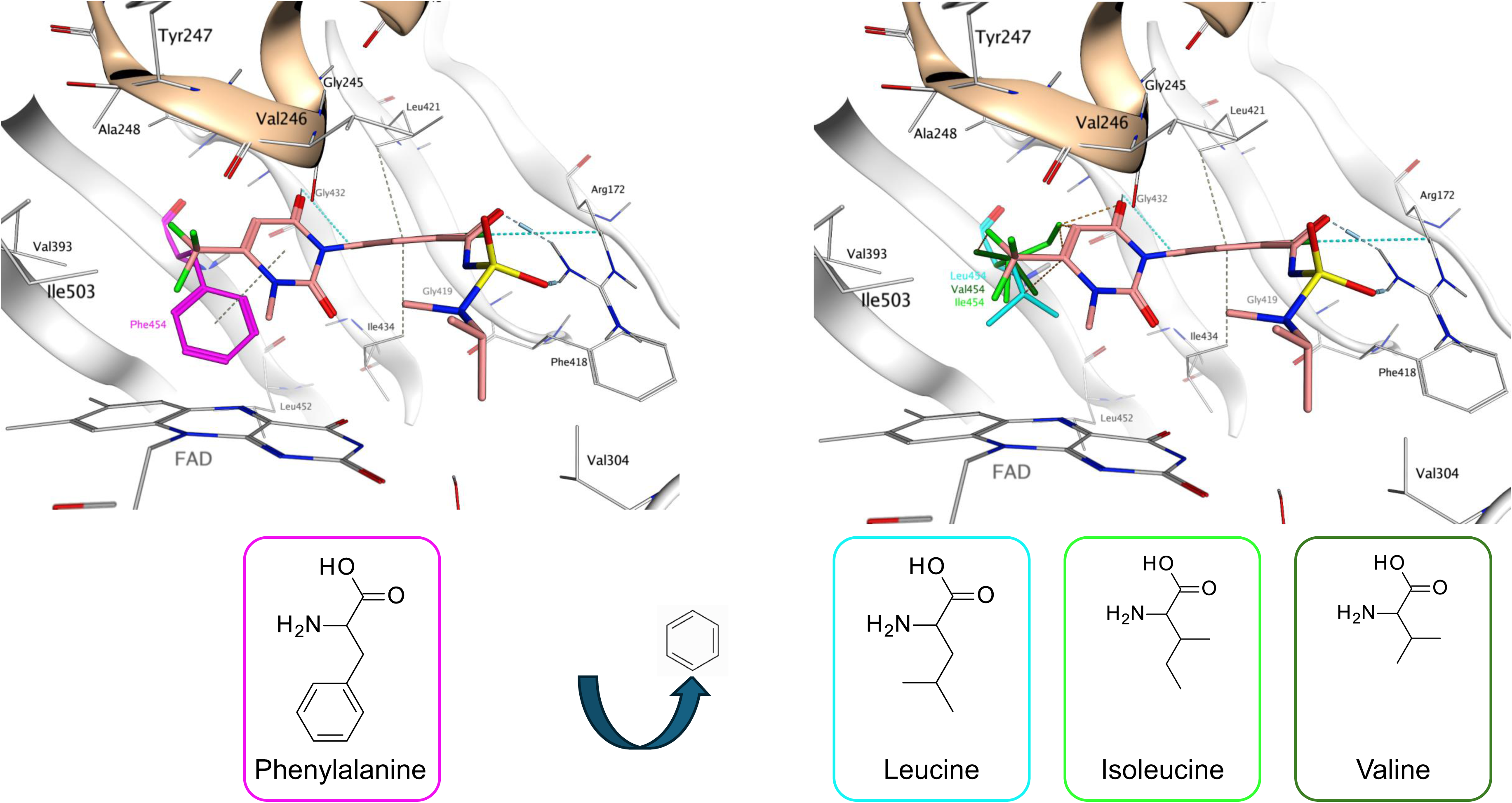
Saflufenacil in the B. scoparia PPO1 Homology Model. The secondary structure of the protein is represented in a white cartoon format. The FAD cofactor is depicted with grey lines. Oxygen atoms are shown in red, nitrogen in blue, sulphur in yellow, chlorine in dark green, and fluorine in light green. Saflufenacil is illustrated in stick form with red carbon atoms. On the left: the uracil warhead of Saflufenacil is positioned between Phe454 (magenta carbon atoms) and Gly245 from the alpha helix (gold), promoting a favorable stacking pi-pi interaction (green dashed line) between Phe454 and the warhead. On the right: substituting Phe454 with Valine (dark green), Isoleucine (green), or Leucine (cyan) eliminates this stacking pi-pi interaction with the phenyl ring side chain of Phe454, and the larger side chains hinder closer binding to the binding site.

Taken together, these findings highlight the structural basis for fomesafen retained efficacy against PPO1 variants harboring F454 substitutions. Its increased conformational flexibility, σ–Π interaction profile, and ability to adjust to altered pocket geometry render it less prone to resistance than carfentrazone-ethyl and saflufenacil. This structural adaptability is a key feature for maintaining herbicidal activity in the presence of target-site substitutions.

### 7. Complementation assays with plant PPO genes in a *hem14* yeast mutant

A PPO herbicide screening system in yeast was developed based on conditional lethality using a hem14/ppo knockout. This strain can still grow on glucose as carbon source but not when the non-fermentable carbon sources ethanol and glycerol are used. However, growth on this medium can be restored by complementation with the *B. scoparia* PPO1 wildtype and F454V I and L substitutions to generate a PPO-inhibitor screening system.

Severe growth inhibition was observed when WT *B. scoparia* PPO1 was expressed in yeast and plated on medium supplemented with 500 µM saflufenacil at pH 3 (Fig. 6). The four sectors (four biological replicates) remained almost entirely clear, without punctate colonies, indicating strong inhibition of PPO1 and loss of yeast viability under herbicide selection. In contrast, yeast strains transformed with *B. scoparia* PPO1 constructs encoding the F454V, F454L, or F454I substitutions displayed vigorous growth under the same conditions with the yeast expressing F454V yielding the maximum growth (Fig. 6). All three variants produced dense, confluent layers of colonies across their inoculated sectors. This strong growth response indicates that each of the F454 substitutions substantially reduces the ability of saflufenacil to inhibit the *B. scoparia* PPO1 variants in yeast.

**Fig. 6.**
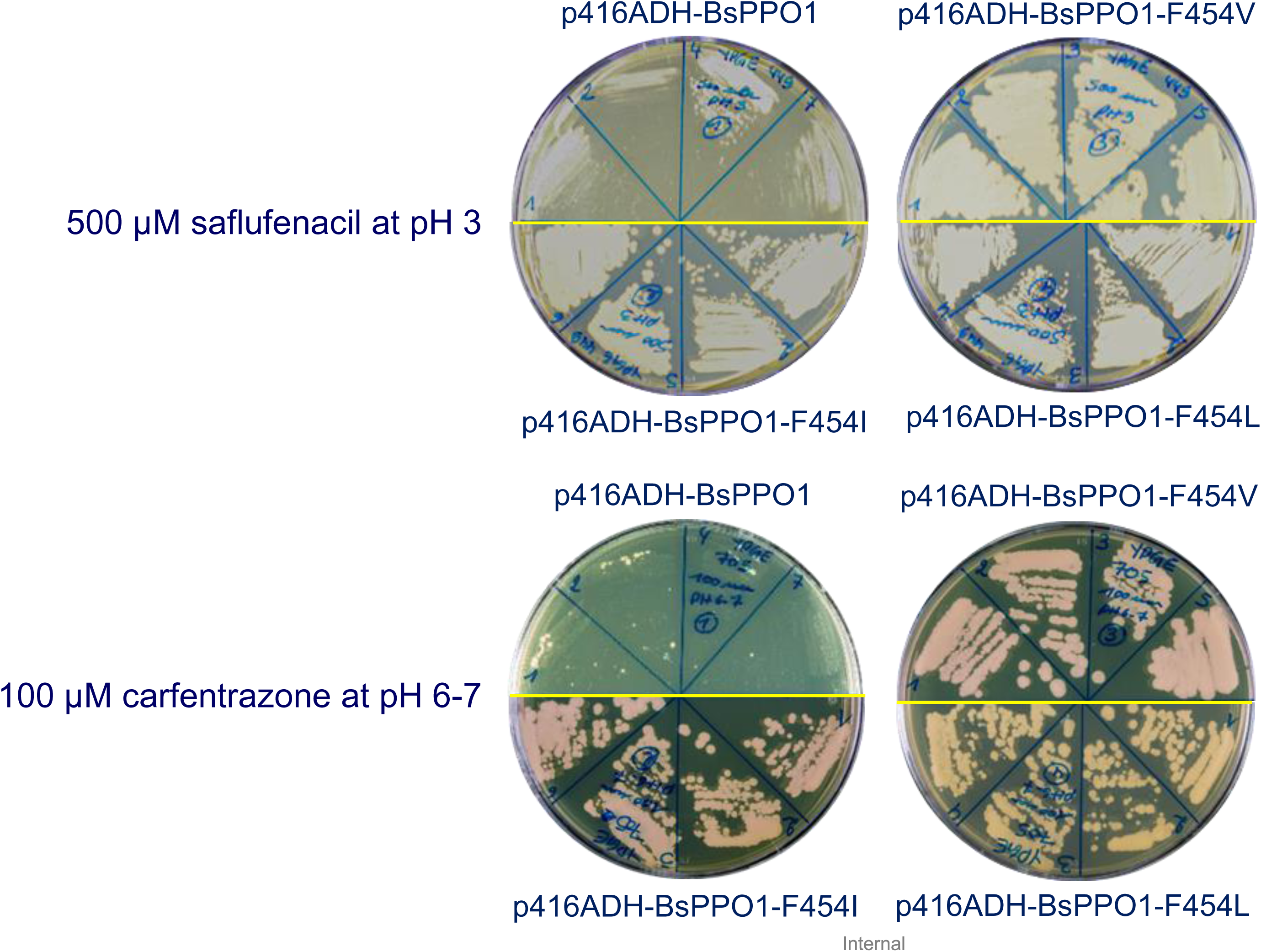
Analysis of p416ADH-BsPPO1 wild-type and mutant strains on Saflufenacil and Carfentrazone. Four ***B. scoparia*** BsPPO1 constructs, wild type and with mutations F454I, F454V and F454L, were screened, each with four biological replicates (upper or lower half of each Petri plate) on 500 µM Saflufenacil (pH 3) and 100µm Carfentrazone (pH6-7), respectively.

When carfentrazone was tested at pH 6–7, yeast expressing the *B. scoparia* PPO1 WT again showed no detectable colony formation, with completely clear sectors (Fig. 6). In contrast, F454V, F454L, and F454I induced abundant yeast growth, with sectors fully covered by large, cream-colored colonies, and yeast expressing F454V yielded the maximum growth as observed for saflufenacil. The consistent growth advantage of all three mutant variants across two chemically distinct PPO inhibitors demonstrates that substitutions at residue F454 confer cross-resistance to both saflufenacil and carfentrazone-ethyl. Thus, F454 substitutions are sufficient to disrupt effective herbicide binding while retaining enzyme functionality in the heterologous yeast expression system. When fomesafen was tested, no significant growth inhibition was observed at 500 µM in any of the yeast complemented strains tested, independently of the medium pH (Fig S2A and B)., This suggests that fomesafen is not efficiently taken up or metabolized by the yeast in the tested conditions.

### 8. Expression pattern of PPO1 and PPO2 during *A. palmeri* and *B. scoparia* early development

To investigate the expression dynamics of PPO1 and PPO2 during early development of *B. scoparia*, transcript levels were quantified at germination and three shoot growth stages, 2.4 cm, 6.3 cm, and 10 cm. Expression levels were normalized to actin and are presented as fold changes (Fig. 7A). At the germination stage, both PPO1 and PPO2 exhibited moderate expression, with fold-change values of approximately 1.45 and 1.40, respectively. This suggests that both genes are transcriptionally active during early seedling emergence, supporting initial plastid development and tetrapyrrole biosynthesis. At the 2.4 cm stage, PPO1 expression remained relatively stable at ∼1.35-fold, while PPO2 showed a slight increase to approximately 2.0-fold. Notably, from this point forward, both genes demonstrated a marked and progressive increase in expression levels, coinciding with active shoot elongation. By the 6.3 cm stage, PPO1 expression rose to ∼2.3-fold and PPO2 to ∼3.5-fold. This trend continued into the 10 cm stage, where PPO1 reached ∼3.3-fold and PPO2 peaked at 4.6-fold, representing over a threefold increase relative to germination. Throughout development, PPO2 consistently exhibited slightly higher expression levels than PPO1 from the 2.4 cm stage onward, indicating a potentially more prominent role in later stages of seedling growth. The progressive upregulation of both genes suggests a developmentally regulated demand for tetrapyrrole biosynthesis, likely associated with chlorophyll and heme production as the photosynthetic machinery develops.

**Fig. 7.**
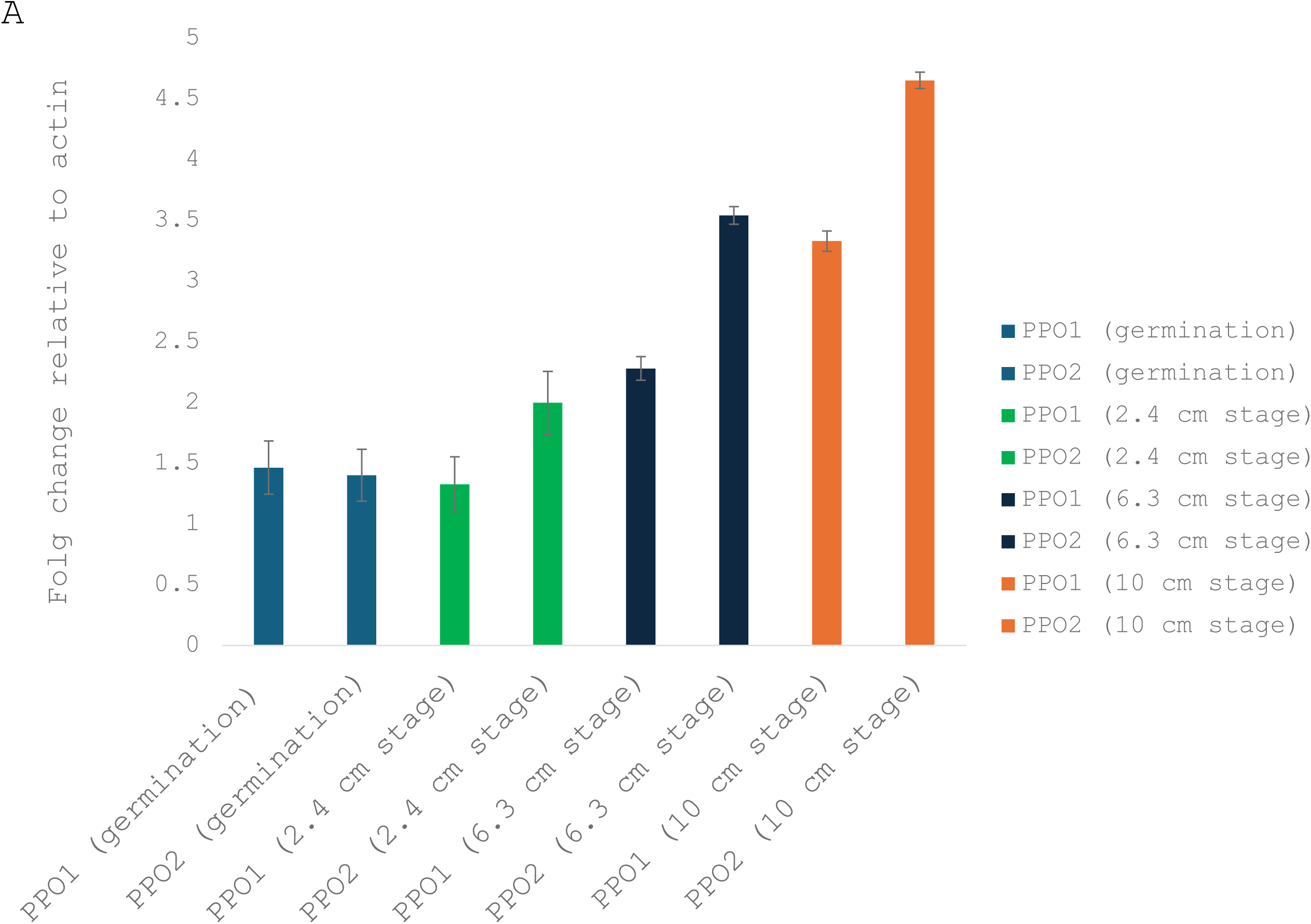
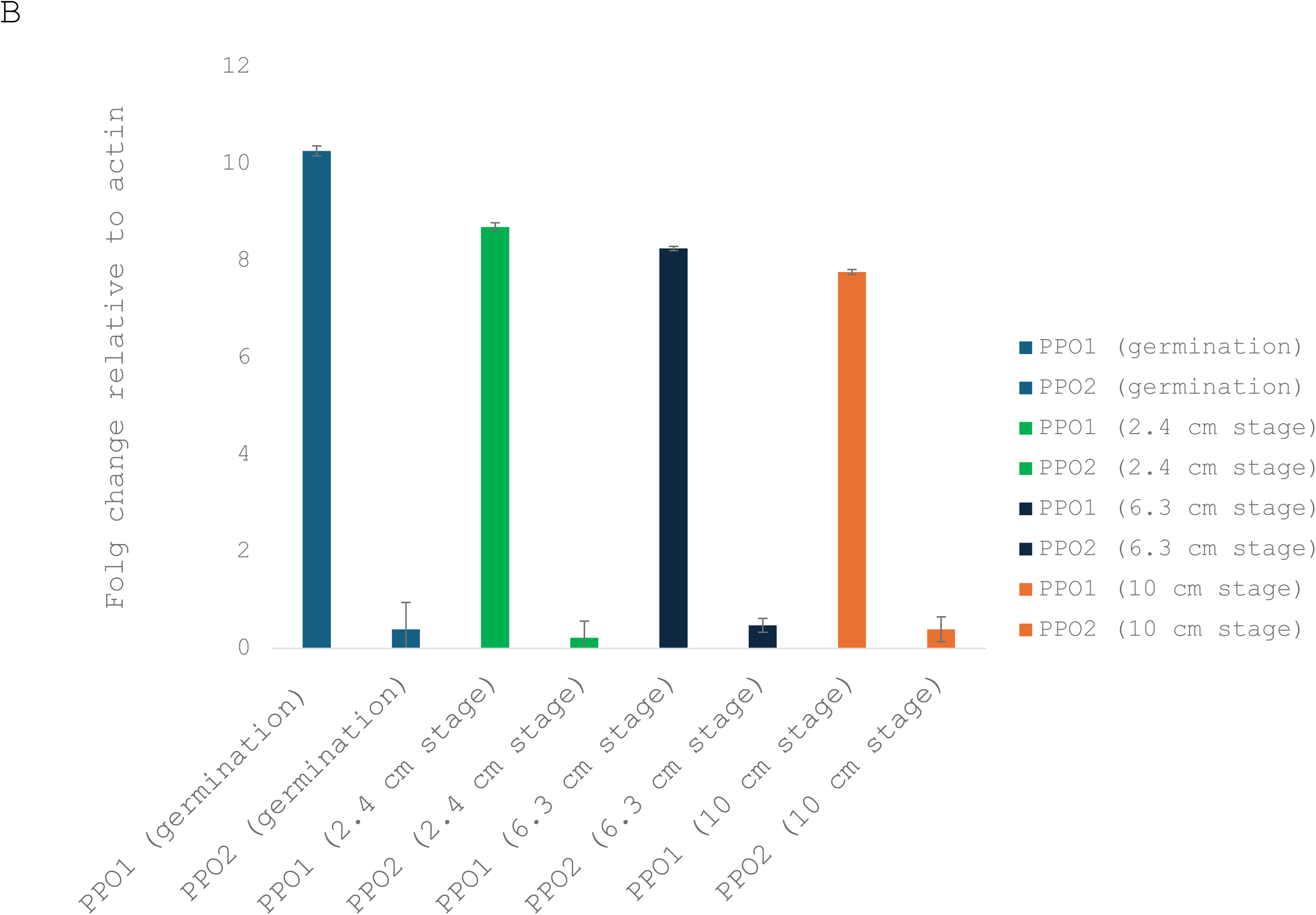
*PPO1* and *PPO2* transcript levels during early seedling development in *B. scoparia* and *Amaranthus palmeri*. Quantitative PCR analysis of *PPO1* and *PPO2* mRNA abundance at four developmental stages in B. scoparia (A) and A. palmeri (B): germination, 2.4 cm shoot length, 6.3 cm shoot length, and 10 cm shoot length using digital PCR. Expression levels are presented as fold change relative to actin. Bars represent 95% confidence intervals (CI95) of three independent biological replicates.

In addition, expression dynamics of PPO1 and PPO2 during early shoot development was also tested in *A. palmeri* at four developmental stages: germination, 2.4 cm, 6.3 cm, and 10 cm shoot length (Fig. 7B). At the germination stage, PPX1 showed very high transcript abundance, with a fold change of approximately 10.5, suggesting a strong transcriptional requirement for PPO1 activity. In contrast, PPO2 expression was much lower, below 0.5-fold, suggesting that the PPO2 isoform might play a minor role during seedling emergence. As seedlings elongated to 2.4 cm, PPO1 expression decreased moderately but remained high (∼8.7-fold), while PPO2 remained very low (∼0.3-fold), with no significant induction. This pattern continued into the 6.3 cm and 10 cm stages, where PPO1 transcript levels remained highly expressed at around 8.2 and 7.8-fold, respectively. In contrast, PPO2 stayed consistently low, with expression values not exceeding 0.4-fold at any stage. These results demonstrate a striking dominance of PPO1 expression across all developmental stages in *A. palmeri*, while PPO2 remains much lower expressed. These data indicate that PPO1 appears to be the primary active isoform throughout seedling development, likely responsible for maintaining plastidic tetrapyrrole biosynthesis needed for chlorophyll production and photosynthetic capacity.

## 4 Discussion

Protoporphyrinogen oxidase–inhibiting herbicides remain an important component of weed management programs; however, resistance to this site of action has expanded rapidly in recent years (Heap, 2025). In dicotyledonous weeds, most confirmed cases of PPO-inhibitor resistance have been associated with target-site mutations in PPO2, particularly the ΔG210 deletion and substitutions at nearby residues (Patzoldt et al., 2006; Salas et al., 2016; Barker et al., 2023). In contrast, we identify here a distinct resistance mechanism in *B. scoparia* mediated by amino-acid substitutions at F454 in PPO1, resulting in strong resistance to saflufenacil and carfentrazone-ethyl while leaving fomesafen efficacy largely unaffected.

### A PPO1-centered target-site resistance mechanism in *B. scoparia*

The resistant *B. scoparia* populations evaluated in this study exhibited high levels of resistance to saflufenacil and carfentrazone-ethyl but remained susceptible to fomesafen. Such herbicide-specific resistance patterns differ from many previously characterized PPO inhibitor-resistant phenotypes, which often show cross-resistance among chemically diverse PPO-inhibiting herbicides (Salas et al., 2016; Rangani et al., 2019; Barker et al., 2023; Riechers et al., 2024; Heap, 2025). Sequencing revealed no known resistance-conferring substitutions in PPO2, and quantitative analyses detected neither CNV nor constitutive overexpression of PPO1 or PPO2, mechanisms that have been implicated in resistance to other herbicide sites of action (Gaines et al., 2010; Jugulam et al., 2014).

Instead, resistance was consistently associated with substitutions at phenylalanine 454 (F454) in PPO1, occurring as F454I, F454L, or F454V. The recurrence of mutations at this residue across resistant individuals strongly suggests directional selection acting on a critical functional site. Although PPO1 has historically been considered a less common target of resistance evolution than PPO2 (Barker et al., 2023), PPO1-based target-site resistance has been documented previously, notably in *Eleusine indica*, where an A212T substitution in chloroplastic PPO1 confers resistance to oxadiazon (Bi et al., 2020). Our findings report the first ever PPO1 target-site mutations endowing herbicide resistance in a broadleaf species.

### Functional validation confirms a causal role for PPO1 F454 substitutions

A defining feature of this study is the integration of biochemical, genetic, and whole-plant evidence to validate causality. Recombinant PPO1 enzymes carrying F454 substitutions exhibited markedly reduced sensitivity to saflufenacil and carfentrazone-ethyl, as indicated by elevated IC₅₀ values, while sensitivity to fomesafen was largely unchanged. Similar enzyme-level shifts have been used to confirm PPO target-site resistance in *Amaranthus* species and other weeds (Patzoldt et al., 2006; Rangani et al., 2019; Barker et al., 2023; Riechers et al., 2024). Transgenic expression of mutant PPO1 alleles in *Arabidopsis thaliana* further confirmed that these substitutions are sufficient to confer resistance at the whole-plant level. Plants expressing PPO1 F454I, F454L, or F454V survived rates of saflufenacil and carfentrazone-ethyl that were lethal to WT controls, while remaining susceptible to fomesafen. These results parallel functional validation approaches used for PPO2 mutations such as ΔG210 and R128 substitutions (Riggins and Tranel, 2012; Salas et al., 2016) and provide strong evidence that resistance in *B. scoparia* arises from altered PPO1 target-site sensitivity rather than non-target-site mechanisms.

### Structural basis of herbicide-specific resistance

Structural modeling suggests that F454 occupies a key position within the PPO1 active site, contributing aromatic and hydrophobic interactions that stabilize binding of certain PPO inhibitors. Replacement of phenylalanine with aliphatic residues is predicted to weaken these interactions, thereby reducing binding affinity for saflufenacil and carfentrazone-ethyl. In contrast, fomesafen appears to rely less heavily on this aromatic contact, consistent with its retained efficacy against F454-substituted PPO1 variants.

These findings align with prior structural and biochemical analyses demonstrating that PPO inhibitors differ substantially in their binding modes and interaction networks within the active site (Koch et al., 2004; Dayan and Duke, 2014). Herbicide-specific resistance resulting from subtle alterations in active-site architecture has also been reported for other PPO mutations and underscores why single amino-acid substitutions do not always produce uniform cross-resistance across the PPO inhibitor classes (Dayan et al., 2018).

### Implications for resistance evolution and weed management

The identification of a PPO1-based resistance mechanism in *B. scoparia* highlights the evolutionary plasticity of this species, which has already evolved resistance to multiple other herbicide sites of action. The herbicide-specific nature of PPO1 F454-mediated resistance suggests that fomesafen may remain effective against populations carrying these alleles, providing a potential management option where agronomically appropriate. Current registered use-cases for fomesafen in North Dakota, USA, and Manitoba, Canada, are limited to preplant, preemergence, or early postemergence in soybean [*Glycine max* (L.) Merr.] or dry bean (*Phaseolus vulgaris* L.) (postemergence in the Red River Valley of Manitoba only), or before emergence of potato in Region 4 of North Dakota (Anonymous, 2025; Ikley et al., 2026). However, *B. scoparia* is not on the label for fomesafen applied alone, and is listed as controlled only when applied with another effective herbicide site of action. Fomesafen is not currently registered for use in Saskatchewan or Alberta, Canada; where PPO inhibitor-resistant B. scoparia is also known to occur (Geddes et al., 2025a; Geddes et al., 2025b). This suggests an opportunity for label expansion of fomesafen in cropping systems of the Northern Great Plains Region. However, extensive reliance on a single PPO inhibitor risks selecting for additional resistance mechanisms, including PPO2-based target-site mutations or non-target-site resistance, both of which have been widely documented in other PPO inhibitor-resistant weeds (Patzoldt et al., 2006; Salas et al., 2016; Dayan et al., 2018; Rangani et al., 2019; Barker et al., 2023; Riechers et al., 2024). From a broader perspective, our results demonstrate that PPO1 represents an underappreciated evolutionary target for resistance and suggest that resistance monitoring efforts should consider variation in both PPO gene family members. Failure to do so may underestimate the diversity of PPO resistance mechanisms emerging in the field.

The evolution of herbicide resistance in *B. scoparia* poses a growing threat to weed management systems throughout the North American Great Plains (Kumar et al., 2019). Resistance to PPO-inhibiting herbicides is particularly concerning due to the broad usage of this site of action (Barker et al., 2023) and its historical efficacy in managing glyphosate-resistant *B. scoparia* populations (Geddes et al., 2025a; Geddes et al., 2025b) that have become widespread in the past decade (Geddes et al., 2022; Geddes et al., 2023; Sharpe et al., 2023; Araujo et al., 2024; Dhanda et al., 2025). For example, PPO-inhibiting herbicides have been identified as important tools targeting glyphosate-resistant *B. scoparia* in fallow (Torbiak et al., 2021a) and prior to growing spring wheat (*Triticum aestivum* L.), canola (*Brassica napus* L.), field pea (*Pisum sativum* L.), and soybean (Yadav et al., 2020; Torbiak et al., 2021b, 2022, 2024), among other crops grown in this region. When PPO inhibitor resistance in *B. scoparia* is considered along with previously-evolved resistance to auxin mimics and inhibitors of ALS, EPSPS, and PSII (specifically, D1 serine 264 binders) (together HRAC Groups 2, 4, 5, 9, and 14), substantial resistance selection pressure will be placed on glufosinate (glutamine synthetase inhibitor; HRAC Group 10), and bromoxynil (PSII inhibitor – D1 histidine 215 binder; HRAC Group 6) alone or in combination with an inhibitor of hydoxyphenyl pyruvate dioxygenase (HRAC Group 27); while other herbicide site-of-action have more-limited use targeting this weed (Geddes et al., 2025a).

The integration of non-chemical tactics and technologies into chemical management programs targeting *B. scoparia* will be necessary given that it has evolved resistance to five herbicide sites-of-action (Kumar et al., 2019; Geddes et al., 2025a), and these traits spread efficiently through pollen- and seed-mediated gene flow (Beckie et al., 2016). Short-lived (1–2 yr) persistence of *B. scoparia* seed in the soil seedbank suggests that densities emerging in-crop are a direct function of seed returned to the soil seedbank the previous fall (Dille et al., 2017; Beckie et al., 2018). Therefore, reducing *B. scoparia* fecundity is critically important to managing this weed efficiently (Geddes and Davis, 2021). In areas with sufficient moisture, fall- or spring-planted cover crops in combination with residual herbicide at termination delay *B. scoparia* emergence and reduce densities in subsequent cash crops (Dhanda and Kumar, 2025). Growing crops with dense, competitive canopies effectively reduced the number of seeds produced by *B. scoparia* plants (Mosqueda et al., 2020). Physical impact mills destroyed viability of *B. scoparia* seed entering the combine during crop harvest (Tidemann et al., 2017; Tidemann et al., 2024). Given the predominant use of PPO-inhibiting herbicides for *B. scoparia* control before crop planting and emergence, strategic and judicious tillage may serve as an alternative non-chemical practice during this weed control window (Osipitan et al., 2019; Obour et al., 2021). Indeed, *B. scoparia* seed persistence was unaffected by burial depth in soil (Dille et al., 2017; Beckie et al., 2018), while emergence declined below 1 cm seed depth and ceased below 8 cm (Schwinghamer and Van Acker, 2008). However, the benefits of tillage for *B. scoparia* management must be balanced against detriments that over-use can have on long-term soil health (Obour et al., 2021). Therefore, continued investment in the discovery of new active ingredients and technologies for *B. scoparia* management before crop emergence will be essential to effectively managing multiple herbicide-resistant populations.

In conclusion, this study provides a comprehensive framework for understanding PPO-inhibitor resistance in *B. scoparia*. The discovery of novel F454 substitutions in PPO1, and their functional validation across multiple systems, underscores the complexity and diversity of resistance mechanisms in weedy species. Our findings emphasize the need for sustained resistance monitoring, integrated weed management, and the strategic use of chemistries like fomesafen that retain efficacy in the face of evolving target-site resistance. The mechanistic insights gained here also inform future herbicide design and provide a valuable model for understanding resistance evolution at the interface of structure, function, and selection.

---

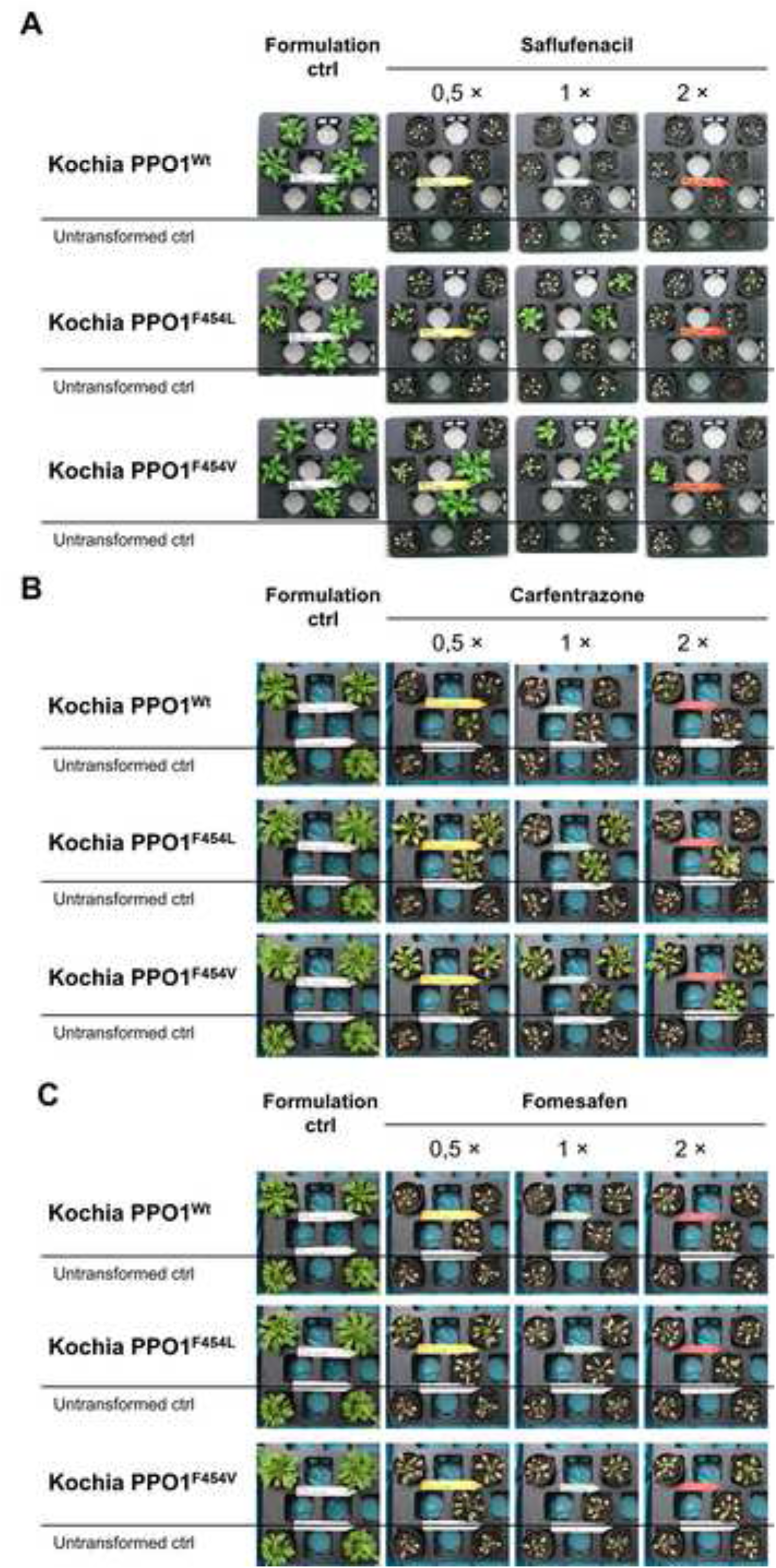

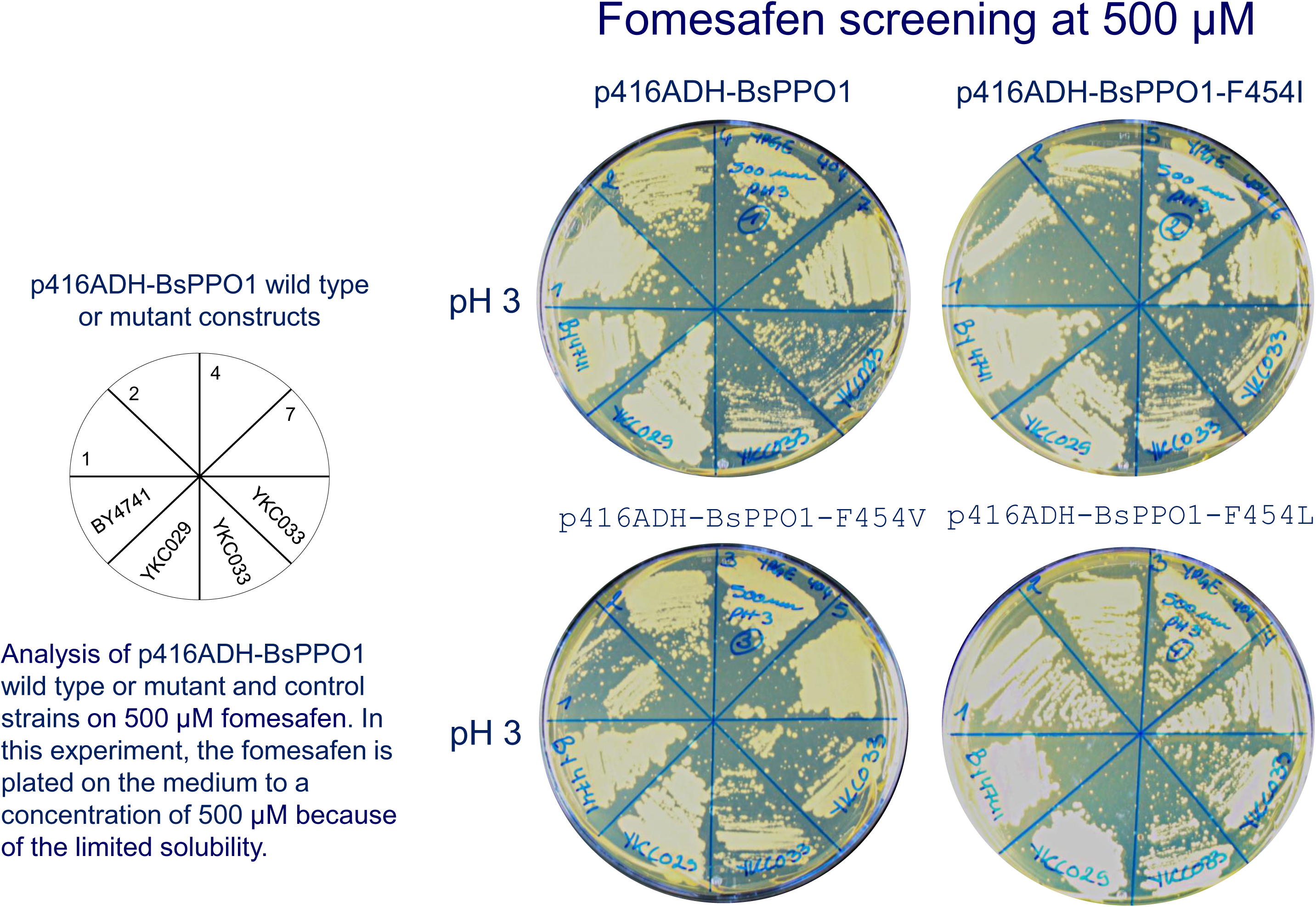
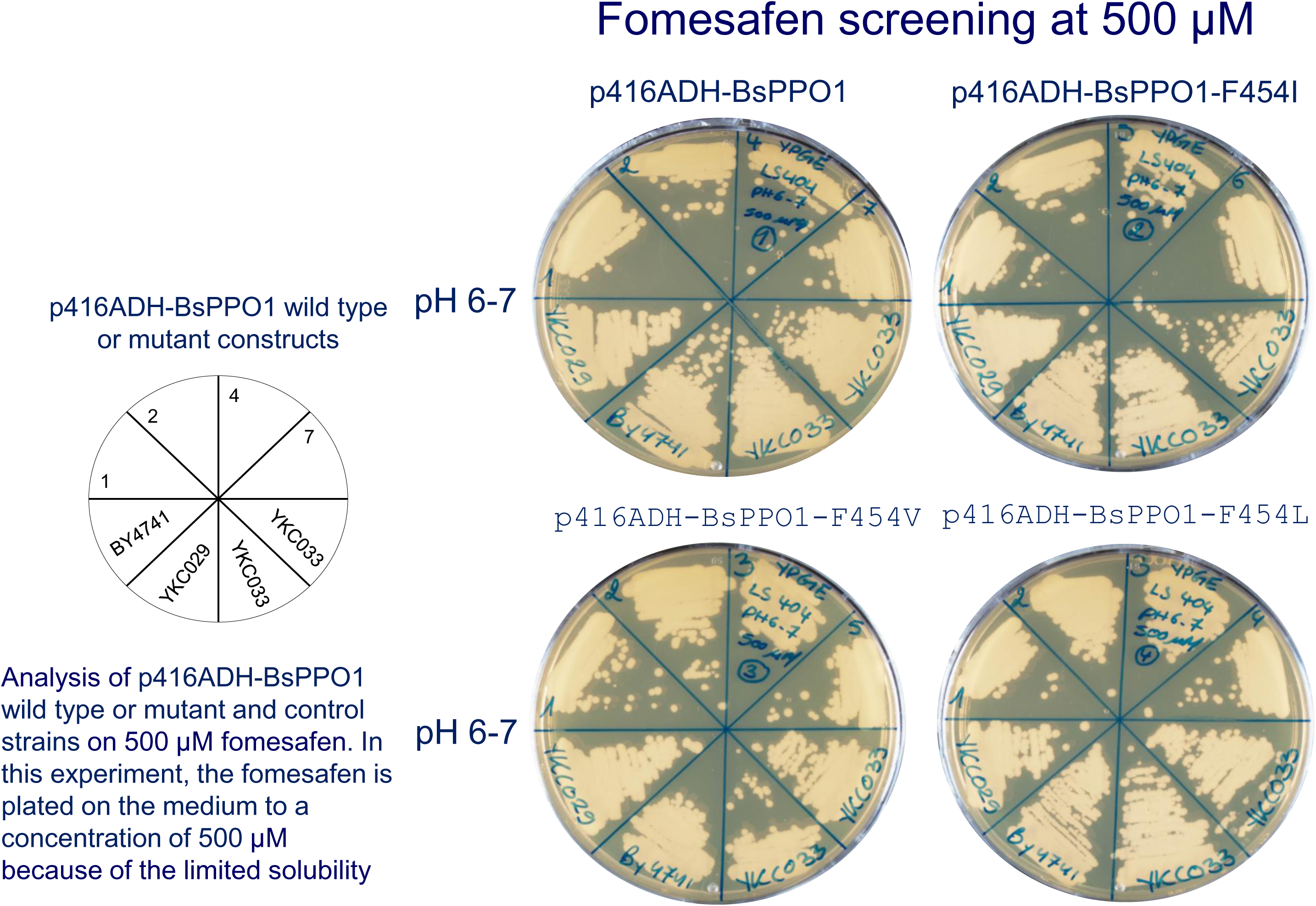

## Notes

### Competing Interest Statement

The authors have declared no competing interest.

